# A scalable hippocampal code for flexible interval timing through persistent activity

**DOI:** 10.64898/2026.04.12.716593

**Authors:** Kori Citrin, Raphael Heldman, Zhuoyang Ye, Yingxue Wang

## Abstract

The brain’s ability to accurately estimate time across various intervals is crucial for flexible behavior. Through the discovery of “time cells”, the hippocampus has emerged as a critical region for temporal processing. However, the hippocampal dynamics underlying flexible interval timing remain elusive. Here, utilizing behavioral perturbations, two-photon calcium imaging, and extracellular recordings in dorsal CA1, we identified a previously unknown pyramidal neuron subpopulation that flexibly scales its activity during interval timing. As mice performed a time-estimation task, CA1 time cells were scarce during the delay period. In contrast, persistently active cells (PACs) exhibited sustained activity from the onset of timing until the animal’s behavioral response. PACs demonstrated temporal scaling: their activity stretched or compressed both with trial-to-trial variations in response time and across different delay durations. PACs comprised two subgroups with complementary ramping dynamics, and their population activity supported reliable decoding of elapsed time. Furthermore, PAC prevalence increased as behavior improved with learning, consistent with their behavioral relevance. Together, these findings revealed scalable sustained dynamics that complement canonical time cell sequences, providing a distinct hippocampal mechanism to support flexible timing behavior.

## Introduction

Accurate time estimation is essential for adaptive behavior. It allows animals to anticipate when reward will become available, suppress premature actions, and initiate motor responses at the appropriate moment. In natural environments, animals must estimate time over a wide range of intervals and adjust their actions accordingly. How neural circuits support flexible time estimation across different intervals while guiding precisely timed actions remains a central open question.

Time estimation is commonly studied under the framework of interval timing, which refers to timing at the level of seconds to minutes. While the frontal cortex and striatum are known to be critical in interval timing behaviors^1–10^, accumulating evidence also suggests a causal role for the hippocampus^11–15^.

Interest in the hippocampus has grown further with the discovery of CA1 “time cells,” neurons that become active at particular moments in time during behaviors that unfold over seconds^16–20^. Together, time cells form a sequence that tiles an interval. This sequential activity is thought to provide a population code for elapsed time. In addition to sequential activity, more recently, hippocampal neurons have been shown to exhibit ramping dynamics that can track time over seconds^21,22^. Thus, multiple representations of time may coexist within hippocampal circuits.

Despite extensive work on hippocampal temporal processing^11–22^, it remains unclear how CA1 neuronal dynamics support time estimation under the variability and flexibility required for behavior. During timing, animals may act earlier or later across trials, and they can adjust their behavior when the required interval becomes shorter or longer. Yet, it remains unclear whether time cell sequences alone support flexible interval timing or whether additional CA1 dynamics contribute.

Here, we combined behavioral perturbations with *in-vivo* two-photon calcium imaging in dorsal CA1 while mice performed a time-estimation task to obtain reward. We found that although time cells were present, they were strongly biased toward the reward and sparsely covered the rest of the interval. In contrast, we identified a subpopulation of CA1 pyramidal neurons, persistently active cells (PACs), that maintained sustained activity from the onset of timing until the initiation of predictive licking. PAC dynamics exhibited temporal scaling when mice responded earlier or later across trials and when the required timing interval became shorter or longer. Furthermore, PACs supported reliable decoding of elapsed time and became more enriched with learning, consistent with a behaviorally relevant role in timing. Together, these results suggest that CA1 can support interval timing not only through sequential activity but through sustained, scalable activity patterns that provide the flexibility required for goal-directed behavior.

## Results

### Reward consumption as a start signal in the time-estimation task

To investigate the hippocampal neuronal dynamics underlying interval timing, we trained mice on a time-estimation task. During the task, head-fixed mice sat in a tube in front of virtual reality screens (Fig. 1a). Each trial began with a 1s visual cue followed by a 4s delay during which the visual scene remained constant. Although mice were not punished for early licking, they received a water reward only if they licked within a 1s unmarked reward window after the delay. Failure to lick within the window resulted in no reward for that trial (Fig. 1b). After training, animals concentrated licking to the period shortly preceding the reward window (“predictive licking”), consistent with estimating elapsed time to anticipate the upcoming reward (Fig. 1c). Trained mice successfully obtained reward on 98.433 ± 0.133% of trials. Local hippocampal inactivation with muscimol significantly impaired predictive licking and reduced the percentage of rewarded trials, confirming that task performance depends on intact hippocampal function (Fig. S1).

**Figure 1:**
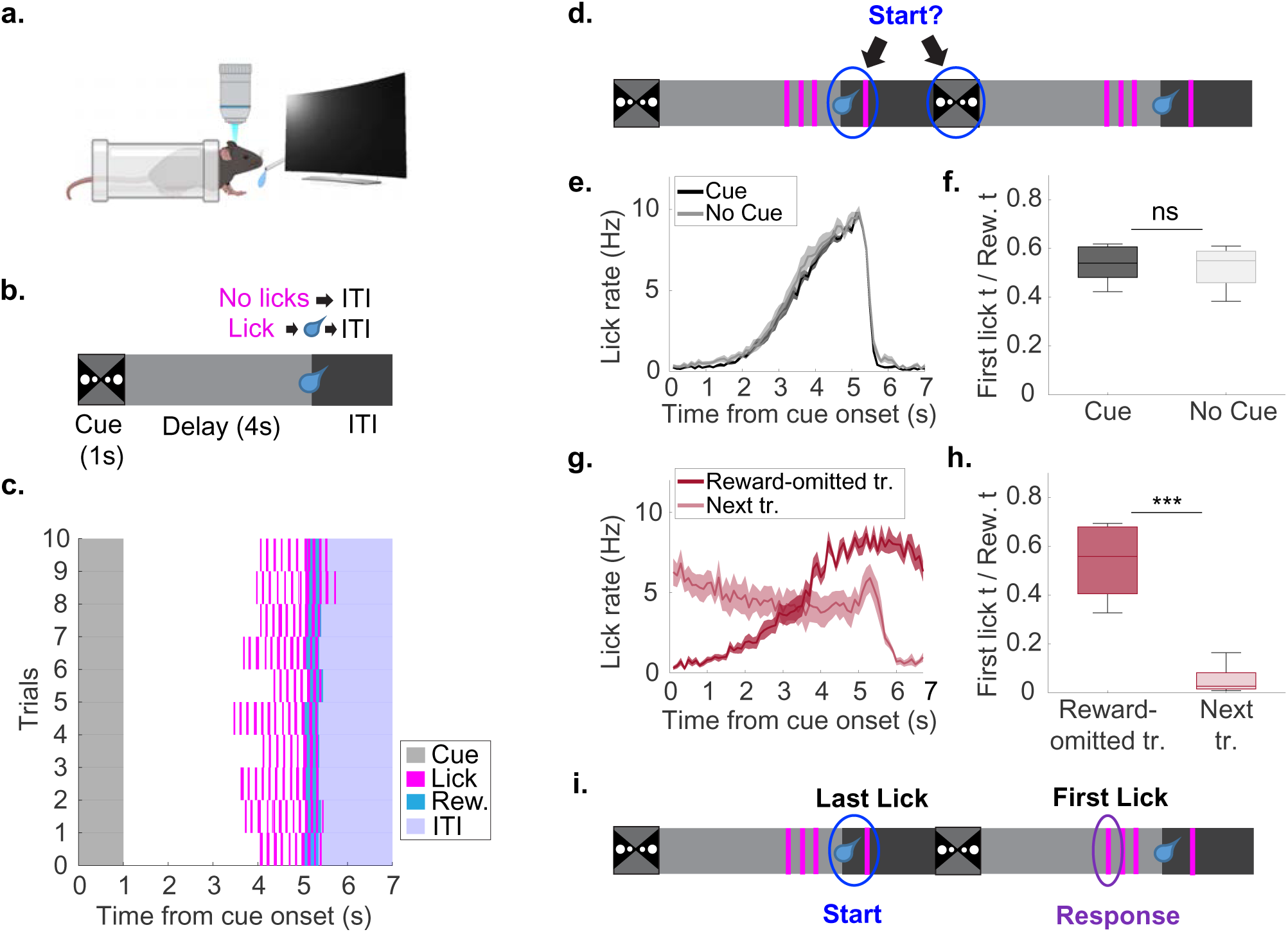
Mice align their timing to the reward consumption event in the time-estimation task. a. Schematic of behavior apparatus. Head-fixed mice sit in a tube in front of virtual reality screens and a lick port. b. Time-estimation task design. After a 1s visual cue and a 4s delay period, mice must lick within a 1s unmarked reward window in order to trigger water delivery. Failure to lick in the reward window results in no reward for that trial. c. Example trials from a well-trained animal. Animals restrict their licking until close to the reward window. d. Schematic of candidate reference points for timing: the visual cue and the reward consumption event. e. Overlaid averaged lick profiles from the cue (black) and no cue sessions (gray). The shaded area represents SEM. f. Predictive licking compared between the cue and no cue conditions. Predictive licking is measured by first lick time / reward time. This provides a measure of how close the animal is to the reward window before it initiates licking (mean ± SEM, cue-present (aligned to cue onset): 0.541 ± 0.032; cue-absent (aligned to typical cue onset time): 0.535 ± 0.034; p = 1.000, Wilcoxon signed-rank test; 3 animals, 6 sessions; mean ± SEM is used unless specifically noted). g. Overlaid averaged lick profiles from the reward-omitted trials and the trials immediately after the reward-omitted trials (“next trials”). h. Predictive licking (first lick time / reward time) compared between the reward-omitted trials and the next trials (reward-omitted trials (aligned to cue onset): 0.539 ± 0.052; next trials (aligned to cue onset): 0.053 ± 0.019; p = 0.008, Wilcoxon signed-rank test; 4 animals, 8 sessions). i. Schematic of the timing behavior in our task: The reward consumption event (reward and last lick on the previous trial) is defined as the “start” of the timing interval, and the first lick on the current trial is defined as the “response”.

In our task, the onset of predictive licking (“first lick”) marks when the animal initiates reward-seeking, providing a trial-to-trial behavioral readout of an animal’s response time. Interpreting the first lick time requires identifying the reference point from which elapsed time is estimated. Because each trial began with a visual cue, we initially hypothesized that the cue onset would serve as this reference point (Fig. 1d). To test if this was the case, we designed behavioral perturbation experiments.

We had mice perform a “no cue session” in which the visual displays were turned off and a “cue session” where visual displays were turned on for within-subjects comparison (Fig. 1e,f; see Methods). To control for the animal’s satiety, we alternated the order of the cue and no-cue sessions across recordings. Removing the cue did not significantly affect predictive licking (Fig. 1e,f; “predictive licking” measured by the time of first lick / reward time), indicating that, once trained, animals do not require the visual cue to perform the task. Therefore, the visual cue is unlikely to serve as the reference signal for timing.

We next asked whether animals timed relative to other salient sensory/motor events, such as the reward delivery (Fig. 1d). In this task, reward delivery and consummatory licking are tightly coupled: the final lick of reward consumption (“last lick”) consistently occurred ∼0.3s after reward delivery (Mean ± SEM: 0.338 ± 0.007s). We therefore treated reward delivery and the last lick as a single reward-consumption event. To test whether this event serves as the reference point for timing, reward was pseudorandomly omitted on 10% of trials. On reward omitted trials, animals initiated licking at similar times as on control trials (first lick time aligned to cue: control trials: 2.845 ± 0.311s; reward omitted trials: 2.864 ± 0.353s; p = 0.844, Wilcoxon signed-rank test, 4 animals, 8 sessions). However, they failed to terminate licking after the expected reward time and continued licking into the subsequent trial. On the subsequent trial, licks were no longer concentrated around the reward window (Fig. 1g-h). This selective disruption of the next trial’s behavior is consistent with the reward-consumption event contributing to the reference signal that initiates the next timing interval.

Based on these results, we define the previous trial’s reward-consumption event as the “start” of the timing interval and the current trial’s first lick as the “response” (Fig. 1i). Unless otherwise noted, going forward, we use the previous trial’s last lick to timestamp the “start”, and “first lick time” refers to the time of the first lick on the current trial relative to this last-lick timestamp.

### Time cell peaks are concentrated around reward consumption and sparsely cover the remainder of the trial

We hypothesized that sequential “time cell” activity in the hippocampus would represent elapsed time during the time-estimation task. To test this, we performed two-photon calcium imaging in dorsal CA1 (Fig. 2a,b). Across sessions, 25.390 ± 1.587 % of CA1 neurons met the time cell criterion (Fig. 2c,d; 1,283/5,161 neurons, activity aligned to the previous trial’s last lick; 11 animals, 15 sessions; see Methods). However, time cell peaks were largely concentrated within the 1s period after the last lick (Fig. 2d,e). This enrichment in the first 1s time bin persisted when identifying time cells aligned to the reward delivery, suggesting that time cells preferentially represent the reward-consumption event (Fig. S2a–b).

**Figure 2.**
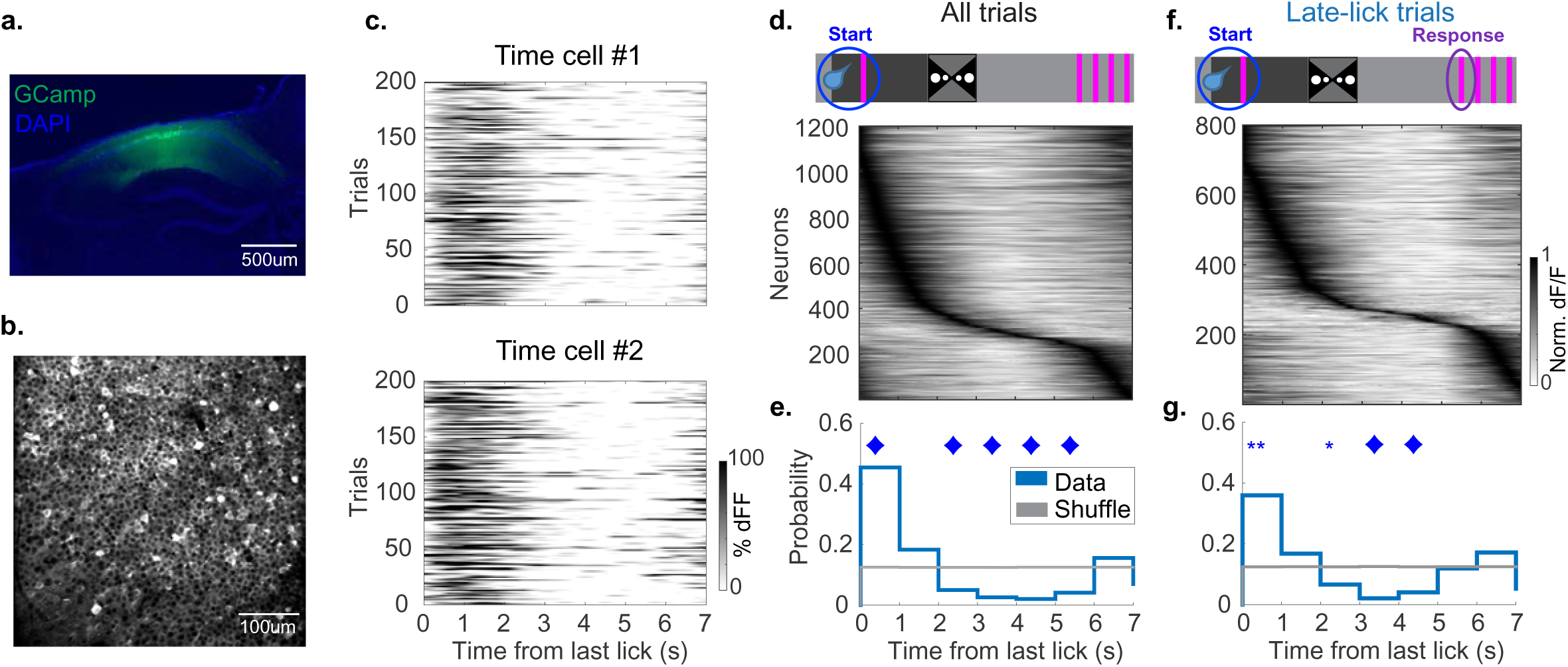
Time cells peaks are biased towards reward consumption and sparsely tile the remainder of the trial. a. Histology of the dorsal hippocampus in a mouse expressing GCamp7f in CA1 (green), with cell nuclei counterstained with DAPI (blue). b. Example field of view (FOV) of the CA1 pyramidal layer in a mouse expressing GCamp7f. c. Heatmap of two example time cells. d. Top: schematic of task. Bottom: heatmap of the time cell sequence aligned to the last lick of the previous trial, including all the time cells detected across all recording sessions. Neurons are sorted by their peak time. e. Histogram of the time cell peaks from 2d (blue) compared to the shuffle distribution (gray) (time bin 0-1s peak probability: real: 0.443 ± 0.037; shuffle: 0.125 ± 0.000; p = 2.329e-05; Diamond: p <0.001, **: p <0.01, *: p <0.05, Wilcoxon rank-sum test; 11 animals, 15 sessions). f. Top: schematic of task. Bottom: heatmap of the time cell sequence aligned to the last lick. Time cells are detected based on activity averaged over the late-lick trials (first lick time 5-6s relative to the last lick). g. Histogram of the time cell peaks from 2f (blue) compared to the shuffle distribution (gray) (time bin 0-1s peak probability: real: 0.347 ± 0.038; shuffle: 0.126 ± 0.001; p = 0.001, Wilcoxon rank-sum test). Time cell activity increases around the time when the first lick occurs (800/5,161 neurons).

In contrast, significantly fewer time cells had peaks in the remainder of the trial, specifically >2s after the last lick. To determine if this non-uniform coverage was due to behavioral variability, we restricted the analysis to trials with similar first lick times (first lick time between 5 and 6s, “late-lick trials”). This analysis yielded a comparable bias in the distribution of time cell peaks (Fig. 2f-g). Furthermore, the analysis revealed that while time cells were sparse during the few seconds leading up to the animal’s licking, many started to become active around the first lick time (“response”) (in this case, 5-6s; Fig. 2f-g).

Thus, although CA1 time cells were present in this task, their peak times were concentrated around the reward consumption event and became sparse during the subsequent waiting period before animals initiated licking.

### Persistently active cells span the interval from start-to-response

We next asked whether additional hippocampal dynamics may represent the period when time cells are scarce. To this end, we quantified pyramidal neuron dynamics aligned to the previous trial’s last lick (“start”) (Fig. S3a; see Methods). This analysis revealed a prominent subpopulation of neurons whose trial-averaged activity increased shortly after the last lick and remained elevated for several seconds (Fig. S3a; dFF_aft_/dFF_bef_ >1.5; see Methods). Notably, 58.766 ± 5.170% of neurons that showed increased activity to the last lick also decreased their activity to the first lick in the current trial (“response”) (Fig. S3b; dFF_aft_/dFF_bef_ < 0.667). Among these neurons, 92.398 ± 1.154% were above baseline for more than three-quarters of the “last-lick→first-lick” interval (baseline defined as the mean amplitude [-1-0]s before last lick), supporting the interpretation that they were persistently active. We therefore refer to this subpopulation as “persistently active cells” (PACs). PACs were similarly identified when classification was performed based on shuffled data (Fig. S4; see Methods), arguing that this result was not due to an arbitrary selection of thresholds.

PACs made up 19.474 ± 2.191% of the total pyramidal neuron population (Fig. S3b; 989/5,161 neurons, 11 animals, 15 sessions; see Methods). In contrast, neurons with the opposite temporal profile—decreased activity to the last lick and increased activity to the first lick—comprised a significantly smaller percentage of neurons (Fig. S5; “lick-on cells”: 6.942 ± 0.925%, 348/5,161 neurons, lick-on vs PACs: p= 1.739e-04, Wilcoxon rank-sum test; 11 animals, 15 sessions). A small percentage of PACs also met the time cell criteria (14.023 ± 0.481%, 134/989 neurons). To test whether persistent activity was specifically enriched for the “last-lick→first-lick” interval, we quantified neurons that were active during the “cue→first-lick” interval (increased activity to cue and decreased activity to first lick) or the “reward→first-lick” interval (increased activity to reward in the previous trial and decreased activity to the first lick in the current trial). Both groups yielded a significantly lower percentage of neurons than the “last-lick→first-lick” interval, suggesting that persistent activity is preferentially enriched during this period (Fig. S6).

Taken together, we identified a subpopulation of neurons, PACs, that increase their activity to the last lick (“start”) and decrease their activity to the first lick (“response”), thus spanning the “start-to-response” interval. This PAC subpopulation covers the period when time cell activity is sparse.

### PAC activity scales with trial-to-trial first lick time

Given that trial-averaged PAC activity decreased around the first lick, we anticipated that PACs would exhibit temporal scaling with first lick time across trials (Fig. 3a). To test this, we separated trials into early- versus late-first lick groups (“early-lick trials”: first lick time between 3.5 and 4.5s; “late-lick trials”: first lick time between 5 and 6s) and compared PAC activity. On late-lick trials, PAC activity remained elevated for longer (Fig. 3b,c). Correspondingly, the temporal center of mass of the activity trace shifted later compared with early-lick trials (Fig. 3c,e (top)).

**Figure 3:**
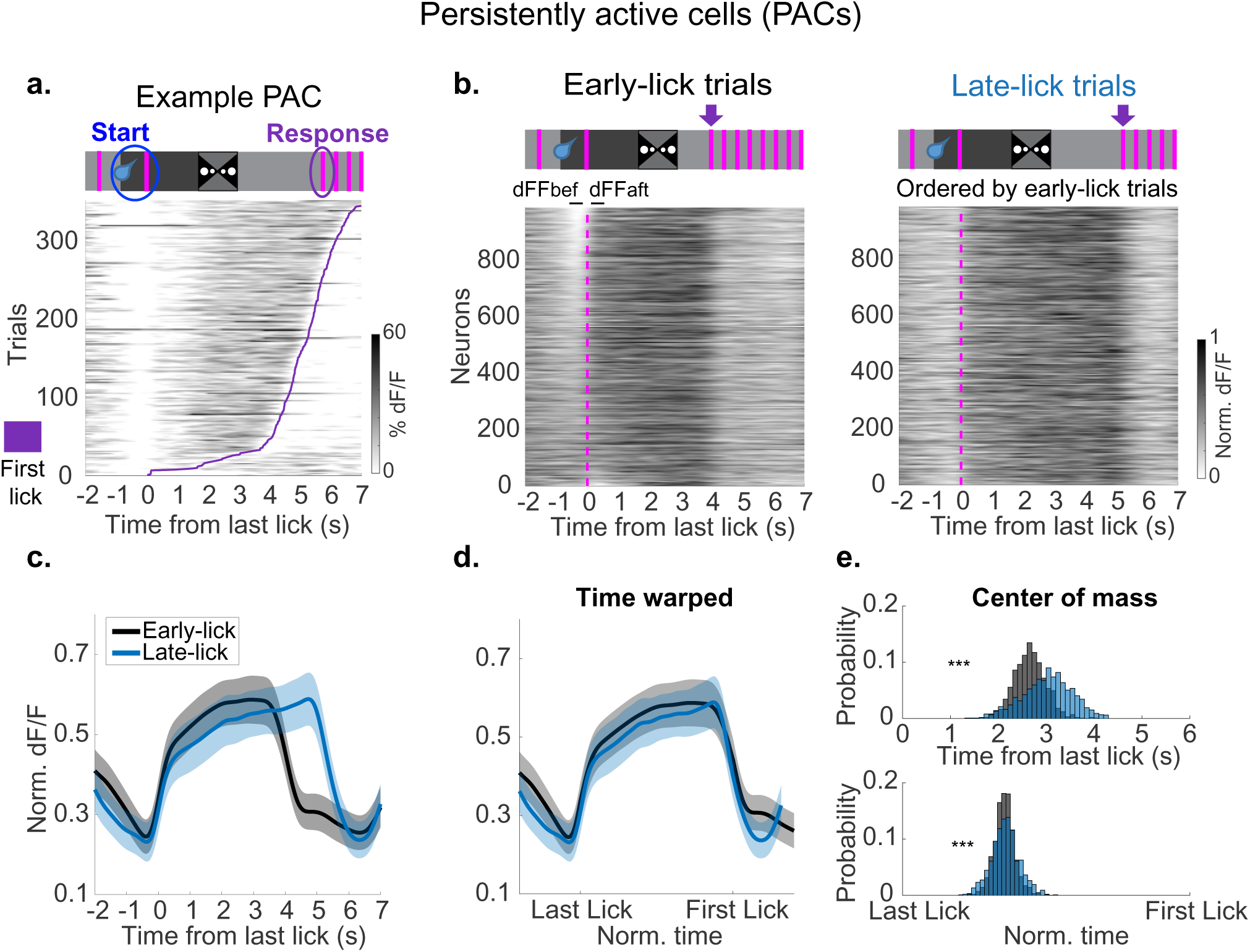
Persistently active cells span the “start-to-response” interval and scale with first lick time. a. Top: schematic of task. Bottom: example persistently active cell. Each row is one trial. Trials are ordered based on the animal’s first lick time (purple line). This neuron increases its activity at the last lick of the previous trial (“start”) and has sustained activity until the first lick of the next trial (“response”). b. Top left and right: schematic of task. Purple arrow points to the first lick. Bottom left: all persistently active cells on trials where the animals’ first lick time is between 3.5-4.5s (“early-lick trials”). Neurons are ordered based on the strength of their response to the last lick (dFFaft/dFFbef) (see Methods). Bottom right: same neurons in the same order but for trials where the animals’ first lick time is 5-6s (“late-lick trials”). Vertical pink dashed line labels the last lick time. c. Averaged normalized dF/F traces for the neurons in 3b. Early-lick trials (black), late-lick trials (blue). The shaded area represents SEM. d. Time warped traces from the last lick to the first lick for the data in 3c. e. Histogram of center of mass (COM) for persistently active cells. Top: non-warped data. Bottom: warped data (non-warped data COM: early-lick trials: 2.658 ± 0.087s; late-lick trials: 3.020 ± 0.134s; p = 2.047e-70, Kolmogorov–Smirnov test, mean difference = 0.363, Cohen’s D = 0.828; warped data COM: early-lick trials: 0.355 ± 0.009; late-lick trials: 0.358 ± 0.013; p = 4.796e-05, Kolmogorov–Smirnov test, mean difference = 0.003, Cohen’s D = 0.066). Although the statistics are significant for both warped and non-warped data, the effect size is much smaller for warped data.

When we time-warped each trial by normalizing the “last-lick→first-lick” interval onto a common duration, PAC activity profiles from early- and late-lick trials largely overlapped (Fig. 3d). After time warping, the center of mass difference between the two trial types was largely reduced (Fig. 3e (bottom)). Taken together, PAC activity exhibits temporal scaling**—**neurons stretch or compress their activity with the animal’s trial-to-trial variation in first lick time.

Because GCaMP fluorescence reflects calcium signals, which have slower dynamics than spikes, we tested whether scalable PAC dynamics were also present at the level of action potentials. Extracellular recordings during the same task recapitulated the key observations: a subset of neurons exhibited sustained firing starting at the last lick, and their firing profiles scaled with first lick time (Fig. S7).

### PAC activity temporally scales with delay duration

In natural environments, animals often need to estimate time across a range of durations. Previous literature suggests that when a timing duration changes, behavioral response time scales proportionally^23,24^. To test whether PACs also scale with different delays, we trained mice on a variant of the time-estimation task in which the delay preceding the reward window alternated between 3s and 5s in a block structure (Fig. 4a).

**Figure 4:**
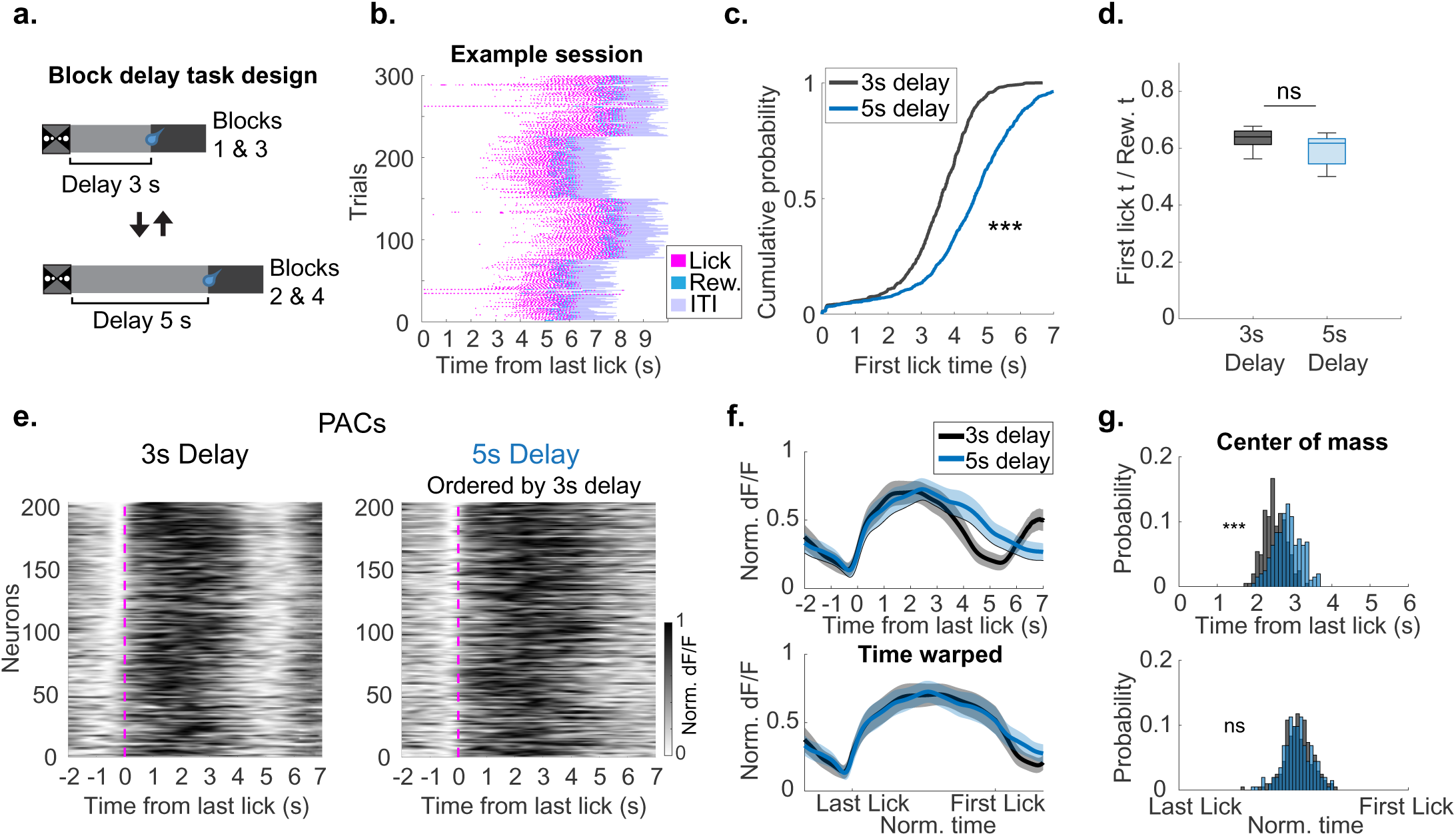
Persistently active cells scale with delay change. a. Schematic of block delay experiment. Blocks 1 and 3 had a 3s delay from the cue offset. Blocks 2 and 4 had a 5s delay from the cue offset. b. Example block delay session behavior, aligned to last lick. c. Cumulative probability of all the first lick times (aligned to last lick) for the 3s and 5s delay blocks (3s delay blocks: 3.439 ± 0.528s; 5s delay blocks: 4.473 ± 0.736s; p = 7.530e-53, Kolmogorov–Smirnov test; 3 animals, 5 sessions). d. Predictive licking measured by first lick time / reward time (aligned to last lick) was not different for 3s vs 5s delay blocks (3s delay blocks: 0.633 ± 0.019; 5s blocks: 0.592 ± 0.028; p =0.151, Wilcoxon rank-sum test; 3 animals, 5 sessions). e. Left: PACs plotted for the 3s delay block trials. Right: Same cells, in the same order but for the 5s delay block trials. PACs that overlap between 3s and 5s blocks are shown. f. Top: Averaged normalized dF/F traces for the neurons in 4e. 3s delay blocks (black), 5s delay blocks (blue). Bottom: time warped activity from the last lick to the first lick for the data in 4f, top. g. Top: PAC center of mass (COM) on the 3s and 5s delay blocks for non-warped data. Bottom: PAC COM on the 3s and 5s delay blocks for warped data (non-warped data COM: 3s delay blocks: 2.510 ± 0.131s; 5s delay blocks: 2.840 ± 0.159s; p = 9.310e-15, Kolmogorov–Smirnov test, mean difference = 0.330, Cohen’s D = 1.014; warped data COM: 3s delay blocks: 0.519 ± 0.028; 5s delay blocks: 0.517 ± 0.028; p = 0.862, Kolmogorov–Smirnov test, mean difference = −0.002, Cohen’s D = −0.029; 204/1,289 neurons).

Consistent with previous studies, animals adjusted their behavior to the delay change: the first lick occurred significantly later in the 5s delay blocks (Fig. 4b,c). However, in the 3s and 5s delay blocks, animals waited a similar fraction of the delay period before licking, consistent with proportional scaling of the animal’s response time^23,24^ (Fig. 4d).

Next, we tested if PAC dynamics altered in parallel with behavior. We first identified PACs separately within individual block types. The percentage of PACs did not significantly differ between the 3s and 5s blocks (% PACs 3s blocks: 26.158 ± 1.290%; % PACs 5s blocks: 20.647 ± 1.740%, p = 0.095 Wilcoxon rank-sum test). Further, there was substantial overlap in PACs across blocks (74.344 ± 1.761%). When focusing on this overlapping population, PAC activity remained elevated for longer during 5s than 3s delay blocks (Fig. 4e,f (top)). Correspondingly, the temporal center of mass of the activity trace shifted later (Fig. 4g (top)). Consistent with lick-time dependent temporal scaling, time warping the “last-lick→first-lick” interval in each block type largely aligned PAC activity profiles (Fig. 4f (bottom)). After time warping, the center of mass difference was no longer significant (Fig. 4g (bottom)).

Our results suggest that PAC activity scales both when animals lick earlier or later from trial-to-trial and when the task delay duration is shortened or lengthened, supporting the idea that PAC activity flexibly tracks different timing intervals. Moreover, PAC dynamics were not unique to immobile behavior. We observed analogous scalable persistent activity in a virtual-reality running task that requires time/distance estimation (Fig. S8; 12.363 ± 3.083%, 464/4,416 neurons; 7 animals, 7 sessions). These results demonstrate that PACs provide a generalizable code for tracking time over different intervals and task conditions.

### Persistently active cells lose their tuning during random licking

One alternative explanation for PAC activity is that these neurons are simply active during lick suppression, rather than being associated with interval timing. To distinguish these possibilities, we recorded an additional “random licking session” (RL) immediately after the time-estimation task. During the RL session, the virtual reality screens were turned off, the delay was increased from 4s to 40s, and water was automatically delivered at the end of each trial. After the dramatic delay increase, animals licked sparsely and sporadically (Fig. S9a; see Methods).

We first asked whether neurons classified as PACs during the time-estimation task retained similar activity during random licking. For a fair comparison, we identified “licking bouts” in the RL sessions (≥5 licks per bout). To match the typical duration of the “last-lick→first-lick” interval in the time-estimation task, we selected pairs of bouts with a 3–6s gap between the last lick of the first bout and the first lick of the next bout (see Methods). Under these matched conditions, neurons classified as PACs in the time-estimation task showed a significantly diminished response to the last lick (Fig. S9b,c). Additionally, a majority of PACs lost their sustained activity during the RL session (RL session: 30.590 ± 5.606% neurons were above baseline for more than three-quarters of the “last-lick→first-lick” interval; Time-estimation task PACs: 89.614 ± 2.769%, p = 6.098-05, paired t-test; 4 animals, 6 sessions, 75 neurons). Therefore, the persistent activity expressed during the time-estimation task was largely absent when animals licked sporadically.

We next asked whether this tuning reduction was due to remapping. That is, whether a different set of neurons became PACs during random licking. To test this, we identified PACs among all the active pyramidal neurons in the matched “last-lick→first-lick” intervals. We detected a significantly lower percentage of PACs during the RL sessions compared with the time-estimation task (Fig. S9d-e). Further, RL session PACs exhibited attenuated responses to the last lick (Fig. S9f).

Together, these results argue against the idea that PACs simply reflect elevated activity during lick suppression and instead support that PACs are preferentially expressed during the time-estimation task.

### Persistently active cells contain decodable temporal information

PACs span the period when time cells are scarce, scale with first lick time, and are more abundant during the time-estimation task than the RL sessions. To further test whether PACs may carry information about elapsed time, we performed Bayesian decoding using PAC population activity alone (see Methods). Time can be decoded at the single-trial level in both the early- and late-lick trials (Fig. 5a). Across sessions, decoding error was significantly lower than a trial-shuffled control for both trial types (Fig. 5b,c; see Methods). These results indicate that PAC population dynamics support reliable decoding of elapsed time from the last lick.

**Figure 5:**
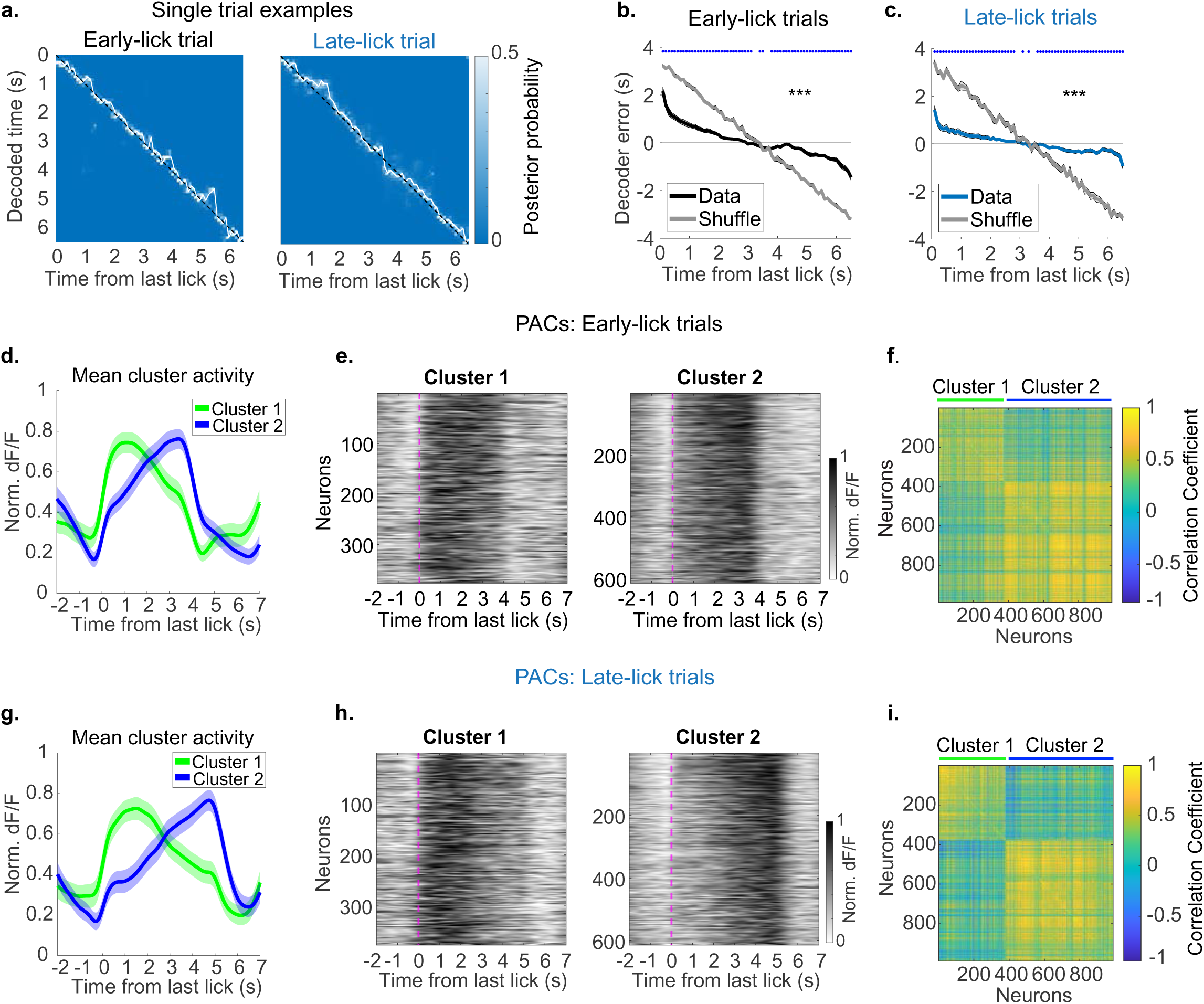
Persistently active cells exhibit ramping activity and can decode time. a. Results from single trial decoding. Left: example early-lick trial. Right: example late-lick trial. Y-axis is decoded time, x-axis is true time. The black diagonal line represents theoretically perfect decoding. The white line is the model prediction. The heatmap represents posterior probability. b. Mean decoder error across sessions for early-lick trials. A decoder error of zero (horizontal line) indicates perfect decoding. The black line represents the model results from the true data, and the gray line represents shuffled data. Time bins with the blue asterisks indicate that the true data is significantly different from the shuffle. The black asterisk represents that the average mean decoder error was significantly lower than shuffle (blue* for each time bin: p <0.05; average mean decoder error: true: 0.521 ± 0.041s; shuffle: 1.632 ± 0.008s, p = 3.392e-06, Wilcoxon rank-sum test; 11 animals, 15 sessions). c. Same as 5b but for late-lick trials. Blue line is the true data, and gray line is the shuffled data (average mean decoder error: true: 0.324 ± 0.050s; shuffle: 1.657 ± 0.024s, p = 3.392e-06, Wilcoxon rank-sum test). d. Results from K-means clustering. Averaged normalized dF/F traces from neurons in cluster 1 (green) and cluster 2 (blue) for early-lick trials. Neurons in cluster 1 peak near the last lick then ramp down to first lick. Neurons in cluster 2 ramp up to first lick. e. Left: heatmap of cluster 1 neurons sorted by the ratio of their response to last lick (dFFaft/dFFbef). Right: Same but for cluster 2 neurons. f. Correlation matrix of neurons grouped by cluster identity for early-lick trials. g-i. Same as 5d-f but for late-lick trials.

To understand how PAC dynamics contribute to this temporal information, we next characterized PAC activity profiles at the single-neuron level^25^. K-means clustering of PAC activity, performed separately for early- and late-lick trials, identified the same two dominant activity patterns in both trial types (Fig. 5d-i, see Methods). Neurons in cluster 1 peaked around the last lick then gradually “ramped down” until the first lick, whereas neurons in cluster 2 gradually “ramped up” to the first lick before shutting down (Fig. 5d,e,g,h). The percentage of neurons within each cluster was similar across trial types (Fig. 5f,i; early-lick trials: cluster 1: 37.715%, 373/989 neurons; cluster 2: 62.285%, 616/989 neurons; late-lick trials: cluster 1: 38.018%, 376/989 neurons; cluster 2: 61.982%, 613/989 neurons). Notably, most neurons maintained their cluster identity across early- and late-lick trials (71.992%; see Methods). Similar ramping patterns were also observed in PACs during the block delay experiment (Fig. S10). Taken together, these results suggest that PACs contain complementary ramp-down and ramp-up subpopulations from which elapsed time can be decoded.

### The percentage of persistently active cells increases with learning

To further investigate the relevance of PACs in interval timing, we asked if PAC prevalence changes over learning. We predicted that PACs would be less prevalent early in training and would increase as timing performance improves over days.

To test this prediction, we performed longitudinal two-photon imaging of dorsal CA1 throughout training on the time-estimation task (Fig. 6a). During the first 3-4 days of training, water was automatically delivered, and the delay was 2s. After day 4, animals had to lick within the unmarked reward window to trigger water delivery. Additionally, the delay started to gradually increase from 2 to 4s. We defined days 1–3 as early-training (2s delay) and days 8–10 as late-training (4s delay).

**Figure 6:**
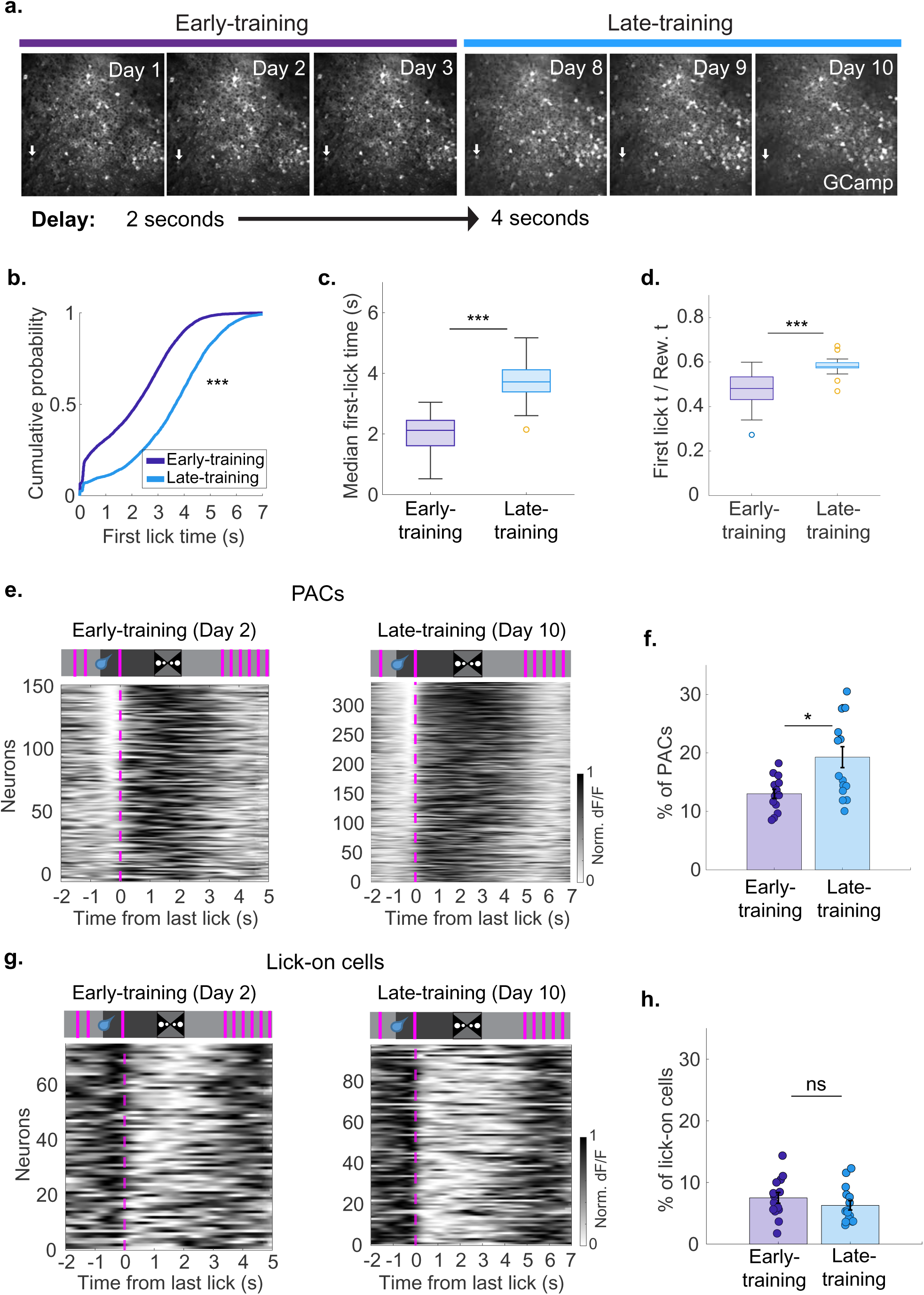
Persistently active cells increase with learning. a. Top: Field of view (FOV) over days for one animal expressing GCamp7f. Days 1-3 are considered early-training and days 8-10 are considered late-training. Bottom: The delay increased from 2s to 4s from early- to late-training. b. Cumulative distribution of all first lick times (aligned to last lick) for early- vs late- training (early-training: 2.077 ± 0.399s; late-training: 3.472 ± 0.437s; p = 5.621e-314, Kolmogorov–Smirnov test; 5 animals, early-training: 14 sessions, late-training: 15 sessions). c. Quantification of the animal’s median first lick time (aligned to last lick) (early-training: 1.923 ± 0.213s; late-training: 3.695 ± 0.193s; p = 2.536e-05, Wilcoxon rank-sum test). d. Predictive licking measured by first lick time / reward time (aligned to last lick) for early- vs late-training (early-training: 0.467 ± 0.024; late-training: 0.582 ± 0.013; p = 4.426e-04, Wilcoxon rank-sum test). e. Left: all PACs across 5 animals on early-training day 2. Right: all PACs from the same 5 animals but on late-training day 10. f. Percentage of PACs increased from early- to late-training. Each point on the scatter plot represents one session (early-training: 13.007 ± 0.778%; late-training: 19.263 ± 1.790%; p = 0.020, Wilcoxon rank-sum test). g. Left: all lick-on cells across 5 animals on early-training day 2. Right: all lick-on cells from the same 5 animals but on late-training day 10. h. Percentage of lick-on cells did not change from early- to late-training (early-training: 7.489 ± 0.874%; late-training: 6.275 ± 0.753%; p = 0.183, Wilcoxon rank-sum test).

Behavior improved significantly across training. Compared with early-training, animals in late-training initiated licking significantly later (Fig. 6b-c). Further, they waited a longer proportion of the delay period before starting to lick, indicating that once trained animals restricted their licks until they were close to the reward window (Fig. 6d).

In parallel with behavioral improvement, PAC percentage significantly increased (Fig. 6e,f). Additionally, the mean trial-to-trial correlation of PAC activity also increased (early-training: mean correlation: 0.006 ± 0.0002; late-training: 0.029 ± 0.001, p = 5.097e-06, Wilcoxon rank-sum test).

A potential explanation for the increase in PACs in late- compared to early-training, is that in early-training, animals often licked continuously, which made the first- and last-lick boundaries ill-defined. To address this, we repeated the analysis after selecting trials with well-defined first and last licks in both training phases (see Methods). Under this restriction, the learning-associated increase in PACs remained robust (Fig. S11a, b).

Finally, we tested whether this learning-related increase was specific to PACs. We quantified the subpopulation with the opposite temporal profile of PACs, “lick-on cells”. In contrast to PACs, the percentage of lick-on cells did not change significantly across training (Fig. 6g,h, S11c,d). Time cell percentage also did not significantly change (early-training: 24.325 ± 4.409%; late-training: 28.051 ± 0.956%; p = 0.132, Wilcoxon rank-sum test, 5 animals, early-training: 14 sessions, late-training: 15 sessions). Thus, learning was more strongly associated with an increase in PAC prevalence compared to other subpopulations. Overall, these results suggest a link between PAC dynamics and improvement in task performance.

## Discussion

In this study, we identified hippocampal population dynamics associated with seconds-scale interval timing. Several main conclusions emerge. First, in a task where animals had to predict upcoming reward based on elapsed time (time-estimation task), CA1 time cells provided sparse coverage of the majority of the time interval before the initiation of predictive licking (“first lick”). Second, we identified a subpopulation of CA1 pyramidal neurons—persistently active cells (PACs)—whose activity increases at the onset of timing and remains elevated until the animal’s first lick, thereby spanning the “start-to-response” interval when time cells are scarce. Third, PAC dynamics exhibited temporal scaling: their activity stretched or compressed both when animals licked earlier or later across trials and when the required delay duration became shorter or longer. Fourth, PAC population activity supported reliable decoding of elapsed time, likely through ramping dynamics. Lastly, the percentage of PACs increased with learning, consistent with their behavioral relevance. Together, these findings suggest that time estimation in this behavioral paradigm is supported by sustained and temporally scaled hippocampal activity, complementing previously reported time cell sequences.

### Time cells are present, but are biased toward reward consumption

Time cells have long been proposed as a hippocampal code for elapsed time, analogous to place cells for spatial position^16–20,26,27^. In this framework, sequential activation across a population of time cells may provide downstream circuits with a readable representation of time^28^. Our data support the presence of CA1 time cells during interval timing, but they also reveal a strong non-uniform distribution: time cell peak activity was scarce during most of the start-to-response interval but was enriched around reward consumption.

One possible explanation for this non-uniform distribution is the limited sensory and motor cues available during the delay in our task. Prior work has shown that place cell and time cell activity are shaped by visual landmarks^29–34^ and movement-related signals^35–40^, such as self-motion and vestibular input. In our task, however, these signals were significantly reduced. After the brief visual cue at trial onset, the visual scene remained constant during the delay. Additionally, animals were head-fixed and largely immobile, thus reducing changing visual, proprioceptive, and vestibular inputs. By contrast, reward consumption was accompanied by strong sensory and motor signals. These signals may contribute to the bias in time cell peaks around the reward, consistent with prior observations that hippocampal activity exhibits reward overrepresentation^34,41,42^. Taken together, the distribution of sensory and motor cues may have favored time cell responses near reward consumption while providing limited representation for the subsequent period preceding the animal’s licking.

A second possibility is that the task requirements may have contributed to the non-uniform distribution of time cells. Previous studies reporting robust time cell sequences often used tasks with stronger working-memory demands, such as alternation paradigms in which animals must retain trial-specific information across a delay^16–20,26^. In contrast, our task required estimating time across a delay period to receive reward but imposed little additional working-memory demand. This is consistent with the possibility that hippocampal temporal coding depends not only on elapsed time itself, but also on the behavioral demands of the task^22,43,44^. Under this view, tasks with stronger memory demands may favor more continuous time cell sequences.

### PAC activity is temporally structured

In our task, the dominant activity pattern spanning the start-to-response interval was not a dense sequence of time fields, but persistent activity in a subpopulation of CA1 neurons. PACs resembled persistent activity described in other brain regions^45–47^. More recently in the hippocampus, Yuan et al. reported that during long delays, reliable time cells were largely confined to the first few seconds, whereas a second population of persistently active cells emerged during the remainder of the delay^48^. This report is in line with our data, suggesting that the hippocampus may represent time intervals using activity patterns beyond canonical time cell sequences. Our findings extend the current understanding by demonstrating that hippocampal persistent activity exhibits flexible scaling with behavior, is correlated with task performance, and carries temporal information.

This temporal information was reflected in complementary ramping activity patterns within the PAC population. One subgroup ramped down toward the first lick, whereas the other ramped up. These ramping dynamics are consistent with ramping signals described in other timing-related circuits - including frontal cortex^3,4,7–9,49^, striatum^5,6,10^, and more recently the hippocampus^21,22,50^-where gradual increases or decreases in activity have been proposed to encode time. Our findings suggest that hippocampal persistent activity is not merely a static delay signal. Instead, it exhibits a sustained yet evolving trajectory across time, combining two features that are often discussed separately: persistent activity and ramping. Together, PACs form temporally structured population dynamics that link the start of an interval to the timing of subsequent action.

### PACs contain time information at the population level

Elapsed time can be decoded from the PAC population activity on single trials. This finding provides a quantitative link between PAC dynamics and time representation. That said, we should interpret this result carefully. Successful decoding does not prove that PACs are the only neural activity pattern relevant for timing, nor does it establish that they are causally necessary for the behavior. Rather, it shows that PAC dynamics contain substantial time information that remains available despite trial-to-trial variation in first lick times. In this sense, PACs provide a population-level signal consistent with a flexible representation of time.

### PAC dynamics show temporal scaling

A central finding of the study is that PAC dynamics showed temporal scaling with first lick time. When animals initiated licking earlier or later across trials, PAC activity compressed or stretched accordingly. PAC activity also scaled when the delay preceding the reward window changed. During 5s delay blocks, mice initiated licking later than on 3s delay blocks, and PAC activity remained elevated for longer in parallel with this shift in behavior. Moreover, time-warping the start-to-response interval largely aligned PAC activity across delay conditions, supporting the interpretation that PACs exhibit temporal scaling.

Temporal scaling has been widely studied in the frontal cortex and striatum, where ramping activity can stretch or compress across different intervals^3–6,8–10^. Our findings extend this idea to the hippocampus. Previous work has shown that hippocampal time cells can scale when interval duration is manipulated^43^, and Yuan et al. reported that persistent activity continues to span the delay as the interval becomes longer^48^. Here, through systematic experiments and analyses, we extend these findings by showing that PACs differ in an important way: they scale not only with imposed delay duration changes, but also with natural trial-to-trial variation in the animal’s response time. Because behavior is not expressed at exactly the same time on every trial, this property suggests that PACs might be suitable to support interval timing under realistic behavioral variability.

### Random licking experiments and learning support the behavioral relevance of PACs

Providing evidence that PAC dynamics are linked to behavior, PAC tuning was markedly reduced during the random licking sessions. This reduction persisted even when analysis was restricted to periods between lick bouts that matched the start-to-response intervals in the time-estimation task. Together with previous work showing that hippocampal representations depend on behavioral engagement^16,51,52^, the loss of PAC tuning suggests that PACs do not simply reflect an absence of licking, but depend on a task structure that requires interval timing.

Further supporting PACs’ behavioral relevance, PAC prevalence increased as task performance improved with learning. Although the delay was longer in the late-training phase, the PAC increase was unlikely to be caused by the delay duration, since PAC percentage was similar in the 3s and 5s delay blocks during the block delay experiment.

The learning related increase was selective: Neither a control subpopulation with the opposite temporal profile of PACs (“lick-on cells”) nor time cells showed a comparable increase in prevalence. This selectivity suggests that the increase in PAC recruitment is not a generic consequence of repeated task exposure but is more specifically associated with the acquisition of the time-estimation task.

One interpretation is that PACs became more strongly recruited as animals learned to concentrate their predictive licking near the reward window. Although the present data do not establish whether PACs are the cause or consequence of improved performance, the close relationship between PAC prevalence and learning supports a link between PAC dynamics and timing behavior.

### Relationship to broader frameworks for hippocampal timing

Our results refine the current views of hippocampal timing in two ways. First, they suggest that seconds-scale timing in CA1 is not necessarily supported by a single population code. Instead, multiple temporal activity patterns may coexist, including time cell sequences and ramp-like persistent activity. In our task, time cells were biased toward reward consumption, whereas PACs spanned the start-to-response interval. This suggests that distinct hippocampal activity patterns may make complementary contributions to timing.

Second, PAC dynamics are consistent with broader frameworks in which elapsed time can be represented not only by a sequence of discrete firing fields, but also by continuously evolving population activity^53^. In this framework, temporal scaling can emerge when the same trajectory is traversed more quickly or slowly under different behavioral conditions. The presence of ramp-up and ramp-down PAC activity patterns together with successful time decoding, suggests that CA1 may represent time through a sustained but dynamically changing population trajectory.

More broadly, these results suggest that the hippocampus may contribute to timing in at least two ways: by representing specific moments via sequential activity, and by maintaining gradually evolving population dynamics that span the interval between behaviorally relevant events.

### Limitations and future directions

Several limitations should be considered. First, the present study identifies PACs as a robust correlational signal linked to interval timing, but does not establish their causal role in timing behavior. Cell-type-specific perturbations or closed-loop disruption of PAC dynamics will be necessary to determine whether they are required for guiding appropriately timed actions.

Second, the circuit mechanisms that generate PAC dynamics remain unknown. PAC activity may arise from temporally structured inhibition, excitatory inputs from the entorhinal cortex or CA3, or neuromodulatory signals associated with reward consumption and anticipation. Dissecting these mechanisms will require simultaneous recordings and targeted manipulations across defined circuit elements.

Finally, the present task reinforces a relatively simple delayed reward-seeking behavior under head-fixation. It will be important to determine how broadly PAC dynamics generalize across more naturalistic settings and other forms of interval timing.

## Conclusion

In summary, our results identify PACs as a hippocampal subpopulation whose activity spans the interval between the onset of timing and the subsequent behavioral response. PAC activity is temporally scaled across various intervals and relevant for task performance during interval timing. These findings suggest that, in this task, CA1 supports interval timing not only through time cell sequences, but through flexible persistent activity that bridges behaviorally relevant events and adapts to changing timing demands.

## Supporting information

SupFigures

## Code availability

Custom scripts for analysis are available on Git-Hub (https://github.com/koricitrin/ManuscriptCode2026)

## Data availability

Data corresponding to all main figures are available at an open-source repository.

## Acknowledgments

We thank D. Fitzpatrick and H. Inagaki for discussions; M. Klement, N. Daniel, and the MPFI machine shop for making mechanical parts for the experimental setups; J. Wells, J. Poole-Capron, H. Shearin, and the MPFI Animal Resources Center (ARC) for animal care. This work was funded by the Max Planck Society, the Max Planck Foundation, and NIH R01 NS119503.

## Author contributions

K.C. and Y.W. conceived the project. K.C., R.H., and Y.W. designed the experiments. K.C. performed all experiments, except R.H. performed electrophysiology recordings and, Z.Y. performed running task experiments. K.C. performed data analysis. K.C. and Y.W. wrote the manuscript with input from all authors. Y.W. acquired funding, provided resources, supervised personnel, and led the project.

## Methods

### Animals

All procedures were performed in accordance with protocols by the Institutional Animal Care and Use Committee at Max Planck Florida Institute for Neuroscience. Both male and female mice (age> 8 weeks) were used in this study. The following mouse lines were used: C57Bl/6J (JAX #000664), SST-IRES-Cre (JAX #13044), Calb2-Cre (JAX #10774), and Ai14 (JAX #007914). Mice were single housed in a 12:12 reverse light cycle and behaviorally tested during the dark phase.

### Surgical procedures

#### Anesthesia and post-operative care

Prior to surgery, mice were anesthetized with 1-2% isoflurane gas. After surgery, mice were administered buprenorphine SR LAB (0.5 mg/kg, SC) and Meloxicam SR (5 mg/kg, SC).

#### Virus injections and viruses

A small craniotomy was performed above dorsal CA1 (relative to bregma, AP: −2.1 mm, ML: −1.7 mm, DV: −1.3 mm). The virus pGP-AAV-syn-jGCaMP7f-WPRE (AAV1) was diluted to 10% of its initial concentration with phosphate-buffered saline (PBS). Then 200 nl of the diluted jGCaMP7f was stereotaxically injected into CA1 using a glass micropipette. Mice used in the running task experiments expressed pGP-AAV-syn-jGCaMP8s-WPRE (AAV9) instead of jGCaMP7f.

#### Hippocampal cranial window and headbar implantation

A 3 mm craniotomy was performed above the left CA1 using a stainless steel circular drill bit. The bone flap was removed with forceps. Then ∼1 mm of cortex was aspirated using a blunt needle connected to a vacuum pump. Chilled saline was used to clear blood and prevent tissue heating. At first, a 20 gauge needle was used to remove most of the cortex. Then, a 25 gauge and finally a 30 gauge needle were used to gently remove the dorsal-most layers of the corpus callosum myelinated fibers^54^. The aspiration was completed once the alveus fibers of the hippocampus were visible. The cranial window implant was composed of a 3 mm glass coverslip, which was attached with UV glue to the bottom of a metal cannula (3 mm in diameter and 1.7 mm deep). The cranial window was inserted into the craniotomy site and then fixed in place with Krazy glue and dental cement. Finally, a stainless steel headplate was attached to the animal’s skull using dental cement.

#### Bilateral cannula implantation for drug infusion experiments

Guide cannulas (26 gauge) were implanted bilaterally above the CA1 region (coordinates: 2.1 mm from bregma, mediolateral ± 1.7 mm from the midline, and 1.1 mm dorsoventral from the brain surface). The cannulas were secured in place with dental cement. Dummy cannulas (33 gauge) of the same length were inserted into the guide cannulas. After the cannula implantation, a plastic headbar was attached to the skull with dental cement, and a wire mesh crown was placed around the headbar to protect the cannulas.

### Behavior and imaging

#### Behavior apparatus & virtual reality system

During the time-estimation task, the animal’s headbar was secured into the head-fixation post with screws, and its body was positioned in a small tube near the head-fixation post. In one setup, the virtual reality (VR) environment consisted of 3 television screens, one directly in front of the animal and two on the sides to account for the mouse’s peripheral vision. On a separate setup, the VR environment was shown on two small screens. A custom software program was written in Unity to display the VR environment.

A microprocessor-based (Arduino) behavioral control system (the miniBCS board, designed at Janelia) interfaced with a custom MATLAB graphical user interface to control the trial structure and water valve. A separate lick port detector (designed at Janelia) was used to convert a touch on a metal lick port into a digital pulse and to send the information to the miniBCS board. Behavioral data were monitored and recorded using MATLAB.

During the running task experiment, the animal ran on a treadmill. Speed was measured using an encoder attached to the back wheel axis. The encoder was also controlled by the Arduino behavioral control system.

#### *In-vivo* 2-photon microscopy

Two-photon imaging was performed using a resonant scanner and a 16x water objective (N16XLWD-PF, 0.8 NA, 3.0 mm WD, Nikon Instruments). A 920 nm excitation laser (Alcor 920 Spark Lasers) was used. Emitted photons were collected using gallium arsenide phosphide (GaAsP) photomultiplier tubes (PMTs) (Hamamatsu). Emission collected was split into red and green channels using bandpass filters (green: FF01-531/46-25 with a range of 508-554 nm, red: FF01-625/90-30-D with a range of 580-670 nm). Images were obtained at 30 Hz sampling frequency under 2x magnification with a 512 x 512 resolution and field of view (FOV) of 477 um x 477 um. Alignment of behavioral events to the imaging signal was done by routing a copy of the imaging start frame signal to the Arduino board used to control the behavior.

The Tdtomato (Ai14) channel in mice was used as a reference to find the same FOV over days.

#### *In-vivo* electrophysiology

Craniotomies were centered around the dorsal CA1 region of the hippocampus using stereotaxic coordinates (AP: −2.1 mm from Bregma; ML: ±1.7 mm from midline). Electrophysiology recordings were performed with a 64-channel silicon probe (Neuronexus, buzsaki64sp). The probe was slowly lowered into the craniotomy site. The CA1 pyramidal layer was identified based on established electrophysiological landmarks, see Heldman et al., 2025^22^, for more details.

#### Drug infusion procedure

Once an animal was trained on the task, it performed one baseline session (pre) of ∼100 trials. After the baseline session, through the bilateral cannula implant we infused of either saline or muscimol. At the beginning of the infusion, the dummy cannula was removed and replaced by an injection cannula, which extended 0.5 mm deeper into the brain than the guide cannula. The injection cannula (33 Gauge) was attached to Tygon tubing (Tygon 720993) with epoxy glue, and the tubing was attached to a 10 μl Hamilton syringe. The syringe was mounted onto a microinjection pump (UMP3 with SYS-Micro4 controller), which slowly injected (100 nl/min) either 200–400 nl of muscimol hydrobromide (Tocris-0289) (1 mg ml^−1^) or saline (0.9%) into CA1. During the infusion, animals performed the time-estimation task with water automatically delivered. Once the infusion was complete, the injection needle was removed, and the dummy cannula was reinserted. One hour after the infusion, mice performed the time-estimation task again for ∼100 trials.

### Behavior paradigms

#### Time-estimation task

Water-restricted mice received 0.6-1.0 ml of water per day. Mice were head-fixed and sat in a tube while performing the time-estimation task. The task structure consisted of a 1s visual cue followed by a delay period with no changing sensory cues. After the delay period, there was a 1s unmarked reward window. Animals had to lick the lick port within the reward window to trigger water delivery; if they did not lick, no reward was given for that trial. Once the reward was delivered or the reward window time had passed, a moving wall signaled the start of the inter-trial-interval (ITI). The ITI randomly varied between 1.5 and 2.5s. Behavioral performance and neural dynamics were comparable in a constant ITI version of the task. After the ITI, the next trial began.

#### Time-estimation task learning protocol

After ∼1 week of water restriction, mice were trained and imaged on the task. For the first 3-4 days of training, the visual cue was followed by a 2s delay period, after which water was automatically delivered. After day 4 of training, the water was switched from automatic delivery to lick-triggered delivery. In the lick-triggered delivery condition, the mice had to lick within the 1s unmarked reward window to trigger the water drop release. Around day 4 of training, the delay length began to gradually increase from 2s to 4s. The delay was increased by 0.5s per day.

#### Time-estimation task with no cue

Mice were trained on the time-estimation task with the virtual reality/visual cues. After learning the task, mice performed one “cue session” and one “no cue session” (140 - 200 trials). During the no cue session, the virtual reality screens were turned off. This eliminated all possible visual cues in the task. The order of the cue session and no cue session were alternated to eliminate the behavioral confound that motivation might change across sessions. The second session was performed immediately after the first session.

#### Time-estimation task with random reward omission

Mice were trained on the time-estimation task with normal reward delivery. After learning the task, they performed sessions with the reward pseudorandomly omitted on 10% of the trials. The pseudorandom reward omission was implemented by the Arduino behavioral control system.

#### Time-estimation task with a block delay structure

A cohort of mice followed the normal time-estimation task training protocol until reaching the 4s delay. After training, they performed a “block delay session”. The first and third blocks had a 3s delay, while the second and fourth blocks had a 5s delay. Each block consisted of 75 trials. There was a total of 300 trials in a session.

#### Running task

Water-restricted mice received 0.6-1.0 ml of water per day. Mice ran head-fixed on a self-paced treadmill. At the beginning of each trial, a visual cue was displayed for 0.5s. Then mice had to run for 180 cm. After the 180 cm, mice had to lick within a 40 cm unmarked reward zone to trigger water delivery. Failure to lick within the reward zone resulted in no water being delivered for that trial. After the water was delivered or the reward zone had past, there was an ITI of 1s where a moving “door stop” appeared to signify the end of the trial. After the ITI, the next trial began.

#### Random licking sessions

Immediately after performing the time-estimation task, mice performed a random licking (RL) session.

During the RL session, the VR screens were turned off, the delay increased from 4s to 40s, and water delivery was switched from lick to automatic delivery. The dramatic increase in the delay length was meant to disrupt the animal’s normal timing process. Since mice performed only 10-20 trials with this new reward contingency, they did not learn the new timing interval, and therefore licked sparsely and sporadically.

Considering that RL trials were dramatically longer than time-estimation task trials, we segmented the RL trials into different licking bouts. A lick bout consisted of a minimum of five licks within 1s. For analysis, we selected pairs of bouts with 3-6s in between the last lick of one bout and the first lick of the next bout to match the “last lick-to-first lick” intervals in the time-estimation task.

### Quantification and Statistical Analysis

#### Two-photon imaging data preprocessing

Tiff files acquired during imaging were uploaded to suite2p^55^ for motion correction, region of interest (ROI) detection, neuropil subtraction, and signal extraction. The suite2p visual interface was then used to manually ensure that the ROIs were pyramidal neurons with a clear signal. Putative interneurons were excluded from analysis.

#### Identification of the same neurons during the time-estimation task and random licking sessions

We identified persistently active cells during the time-estimation task (see below). To identify the same neurons during the random licking (RL) sessions, we used the open-source MATLAB package ROIMatchPub (https://github.com/ransona/ROIMatchPub). We uploaded the suite2p files from the time-estimation task session and its corresponding RL session. ROIMatchPub found overlapping ROIs between the suite2p masks in both sessions. It is important to note that suite2p detects ROIs based on activity. Therefore, we can only compare ROIs that had activity in both the time-estimation task and the RL session. Only 50.790 ± 1.592% of neurons in the time-estimation task were also active in the RL session. Because of this relatively low percentage, we did not do the same analysis with the learning dataset that contains 10 sessions.

#### Time cell classification

Time cells were classified by taking the mean of a neuron’s dF/F across trials. We found the time bin where the neuron’s trial-averaged activity reached its peak, and compared the peak amplitude to a null distribution. The null distribution was generated by circularly shifting the neuron’s activity for each individual trial. We then found the peak amplitude of the mean dF/F across shuffled trials. This was repeated 100 times to build a null distribution of peak amplitudes. A neuron had a significant peak if its peak amplitude was greater than 99% of the null distribution values^26^. In addition to having a significant peak, neurons had to have in-field activity for at least 20% of the trials, and the neuron’s field width could not exceed 4s.

#### Identification of activity-increasing and decreasing neurons

##### Threshold method

We aligned neural activity to various trial events (cue onset, reward delivery, first lick, and last lick). Based on trial-averaged activity, a neuron was considered to have increasing activity to an event if its mean activity amplitude after the event onset (0-0.5s, “dFF_aft_”) divided by its mean activity amplitude before the event onset (−0.5-0s, “dFF_bef_”) was greater than a threshold of 1.5. A neuron was considered to have decreasing activity if its dFF_aft_/dFF_bef_ ratio was less than 0.667.

##### Shuffle method

With shuffling, we determined activity-increasing and decreasing neurons by comparing each neuron’s dFF_aft_/dFF_bef_ ratio to a null distribution of ratios. The null distribution was generated by circularly shifting the neuron’s activity on each trial then finding the dFF_aft_/dFF_bef_ ratio using the trial-averaged dF/F trace. This was repeated 1,000 times to build a null distribution of ratios. A neuron was considered significantly increasing if its ratio was greater than the 90^th^ percentile of the shuffle distribution, while a neuron was considered significantly decreasing if its ratio was lower than the 10^th^ percentile.

#### Persistently active cell classification

A neuron was classified as a persistently active cell (PAC) if its activity increased to the last lick and decreased to the first lick (see Identification of activity-increasing and decreasing neurons). For electrophysiology data, PACs were identified with a FR_bef_ range of −0.5-0s and FR_aft_ range of 0-1s instead of 0-0.5s.

#### Lick-on cell classification

A lick-on cell was a cell whose activity increased to the first lick and decreased to the last lick (see Identification of activity-increasing and decreasing neurons – threshold method).

#### Percentage of the “last-lick→first-lick” interval during which PACs were active

We took the last-lick-aligned trial-averaged activity trace of PACs, and then rescaled the trace so the amplitude ranged from 0 to 1. We calculated the baseline for each cell by looking at the mean activity amplitude in the 1s window preceding the last lick. We then found the time bins where activity was greater than baseline between the last lick and the median first lick time on the next trial. By dividing the number of time bins above baseline by the total number of time bins we calculated the percentage of the “last-lick→first-lick” interval during which the neuron was active. This gave us a measure of the neuron’s sustained activity. We considered a neuron to be “persistent” if its activity was above baseline for greater than 75% of the “last-lick→first-lick” interval.

#### Predictive licking quantification

Predictive licking was quantified by:

First lick time / reward time

This gave us a measurement of how well the animal timed its licks relative to the reward.

A score of zero means the mouse started licking immediately when the trial started, whereas a score of 1 means the mouse’s first lick was at the beginning of the reward window.

We only considered trials where the score was between 0 and 1. This was to exclude trials, during early-training, where water was automatically given and the animal waited to initiate licking until after the water was delivered (i.e., not predictive).

First lick and reward time were relative to the cue onset for Figures 1 and S1 and relative to the last lick of the previous trial for Figures 4 and 6. That said, the results for the comparisons of predictive licking remained the same regardless of whether licks were aligned to the last lick or the cue.

#### Early and late first licks

Early- and late-lick trials are defined by the first lick time on the current trial relative to the last lick time on the previous trial. For the time-estimation task calcium imaging dataset, on early-lick trials, the animals’ first lick time was 3.5-4.5s. On late-lick trials, the first lick time was 5-6s. For the electrophysiology data, on early-lick trials, the first lick was 4.5-5.5s, and on late-lick trials, the first lick was 6-7s (mice from the electrophysiology cohort generally licked later than the imaging cohort). The running task trials are shorter in length than the time-estimation task trials. For the running task, on early-lick trials, the first lick time was 2-3s, and on late-lick trials, the first lick time was 3.5-4.5s.

#### Time warping

We normalized the time axis between the last lick of the previous trial and the first lick of the current trial for each PAC by linearly scaling neuronal activity. FirstLickTimeTarget was set to 1. To scale the activity we used the following:

ActivityTimeWarp = TimeBinsOriginal × (FirstLickTimeTarget/FirstLickTimeActual).

#### Center of mass

The center of mass was calculated by:

COM = sum(time* NormActivity)/ sum(NormActivity)

where NormActivity is the normalized mean dF/F of a neuron and time is a vector of all the time bins.

For the electrophysiology dataset, COM was calculated from 0 to 7s because the late-lick trials were extended to 7s. For all other analyses, COM was calculated between 0 and 6s. For the time-warped data, the number of bins matched that of the non-warped dataset, so the COM calculation was comparable.

#### K-means clustering

We performed K-means clustering on the trial-averaged activity of persistently active cells. We performed clustering separately on early-lick and late-lick trials. The optimal number of clusters was determined by computing the within-cluster sum of squares (WSS/Elbow) and the mean silhouette score.

#### Bayesian decoding

Bayesian decoding of time was performed with the MATLAB function fitcnb. For each imaging session, we trained the model with the activity of persistently active cells on 90% of trials and tested on the other 10% of trials using k-fold cross-validation. The time bin for decoding was 0.1s. Shuffled data was obtained by shuffling the model predicted time for each trial then getting the mean decoder error based on the shuffle.

#### Sub-selection of trials with similar licking behavior

When identifying trials with clear first and last licks, we defined “valid first licks” as trials where the animal waited at least 1s after the cue onset to lick. “Valid last licks” were when the animal stopped licking at least 0.9s before the ITI end. For analysis purposes, we used trials with a valid last lick on the previous trial and a valid first lick on the current trial. This excluded trials where the animals licked continuously across the trial boundary.

