## Supplementary material for "A scalable hippocampal code for flexible interval timing through persistent activity": SupFigures

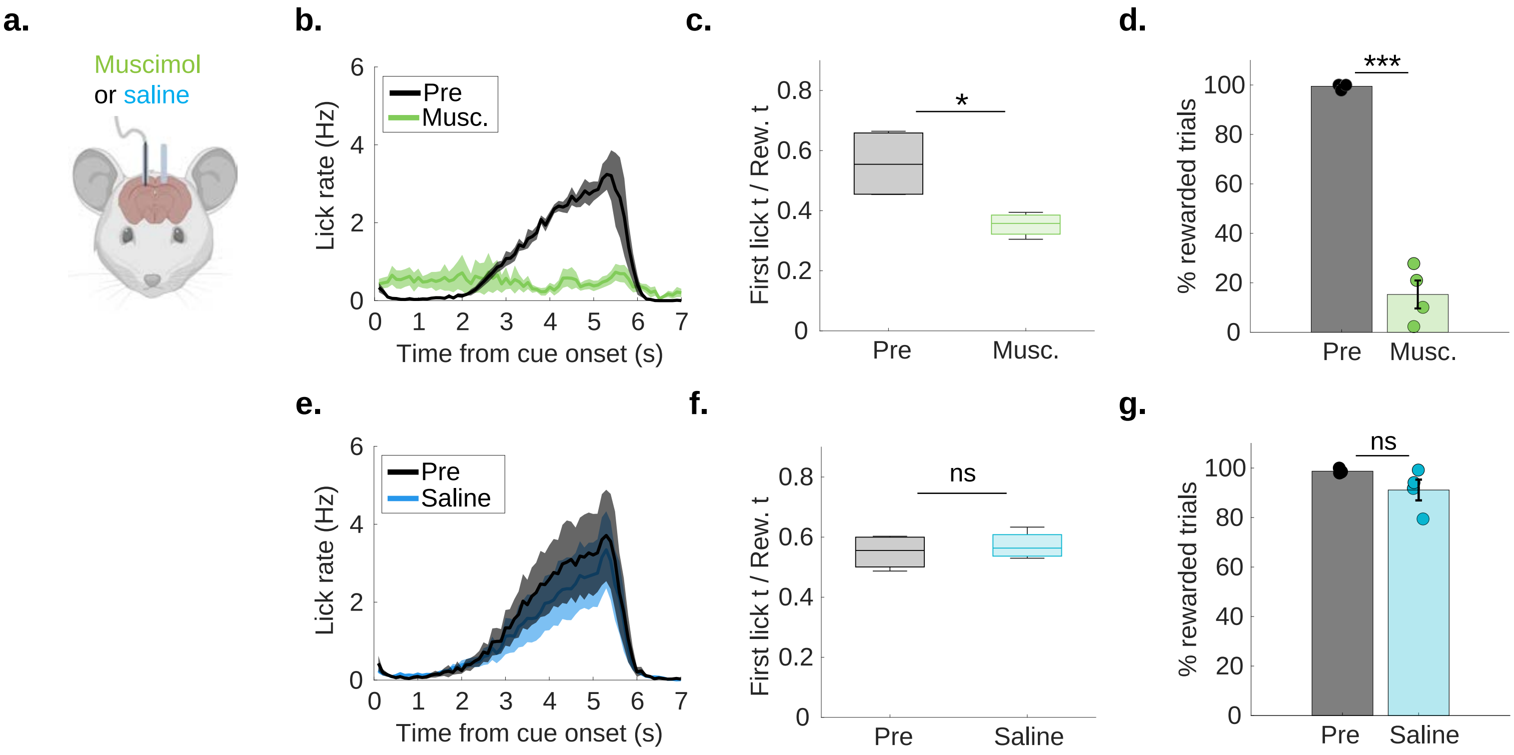


**Supplemental Figure 1:** Time estimation task is hippocampal dependent

a. Schematic of bilateral cannula implant for drug infusion experiments.

b. Averaged lick profiles from the muscimol sessions ("Musc.", green) and pre drug infusion control sessions ("Pre", black).

c. Predictive licking measured by first lick time / reward time (aligned to cue onset) (pre: 0.557 ± 0.059; muscimol: 0.354 ± 0.020; p = 0.035, paired t-test; 4 animals, 4 sessions per condition).

d. Percent rewarded trials (pre: 99.474 ± 0.526%; muscimol: 15.294 ± 5.658%; p = 7.296e-04, paired t-test).

e-g. Same as a-d but for saline. (first lick time / reward time: pre: 0.550 ± 0.029; saline: 0.572 ± 0.023; p = 0.486, paired t-test; percent rewarded trials: pre: 98.716 ± 0.444%; saline: 91.123 ± 4.194%; p = 0.160, paired t-test; 4 animals, 4 sessions per condition).

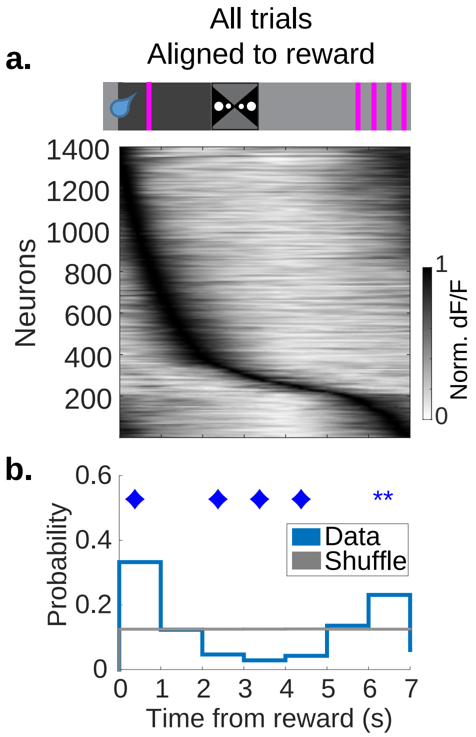


**Supplemental Figure 2:** Time cell sequence aligned to reward

a. Top: schematic of task. Bottom: heatmap of the time cell sequence aligned to reward delivery. Neurons are sorted based on their peak time. Time cells were identified based on activity averaged over all trials aligned to the reward delivery.

b. Histogram of the time cell peaks from S2a (blue) compared to the shuffle distribution (gray) (time bin 0-1s peak probability: real: 0.320 ± 0.030; shuffle: 0.124 ± 0.000; p = 7.792e-04, Diamond: p <0.001 , **: p <0.01, *: p <0.05, Wilcoxon rank-sum test; 1,400/5,161 neurons, 11 animals, 15 sessions).

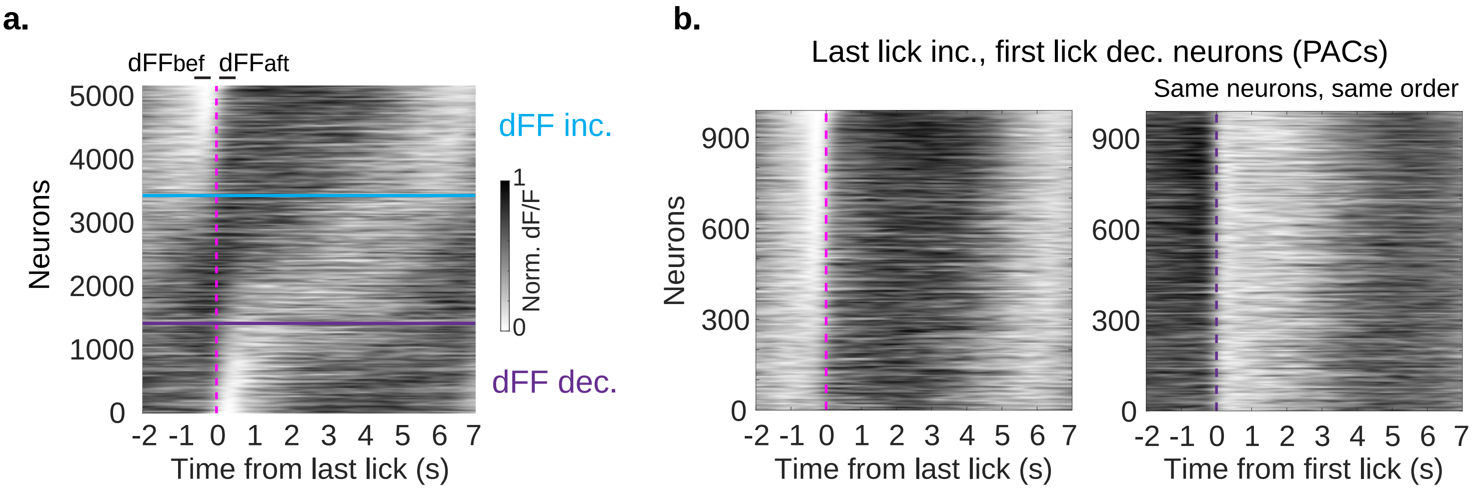


**Supplemental Figure 3:** Pyramidal neuron population aligned to the last lick

a. All recorded CA1 pyramidal neurons aligned to the last lick. Neurons are ordered based on the strength of their response to the last lick ($dFF_{aft}/dFF_{bef}$) (see Methods). Neurons above the blue line increase activity to the last lick ($dFF_{aft}/dFF_{bef} > 1.5$). Neurons below the purple line decrease activity to the last lick ($dFF_{aft}/dFF_{bef} < 0.667$). Neurons between the blue and purple lines were not significantly modulated by the last lick (5,161 neurons, 11 animals, 15 sessions).

b. Persistently active cells ("PACs", neurons that increase their activity to the last lick and decrease their activity to the first lick). Left: heatmap of PACs aligned to the last lick. Neurons are ordered based on the strength of their response to the last lick ($dFF_{aft}/dFF_{bef}$). Right: Same neurons in the same order but aligned to the first lick.

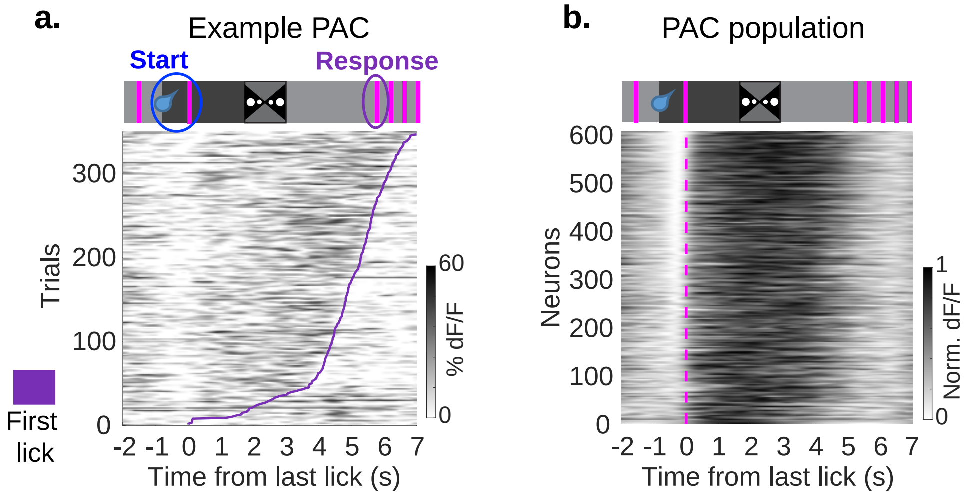


**Supplemental Figure 4:** Persistently active cells identified using a shuffling method

a. Top: schematic of task. Bottom: example persistently active cell identified with a shuffling method. Each row is one trial. Trials are ordered based on the animal's first lick time (purple line).
b. Top: schematic of task. Bottom: all persistently active cells identified with a shuffling method. Neurons are ordered based on the strength of their response to the last lick ($dFF_{aft}/dFF_{bef}$).

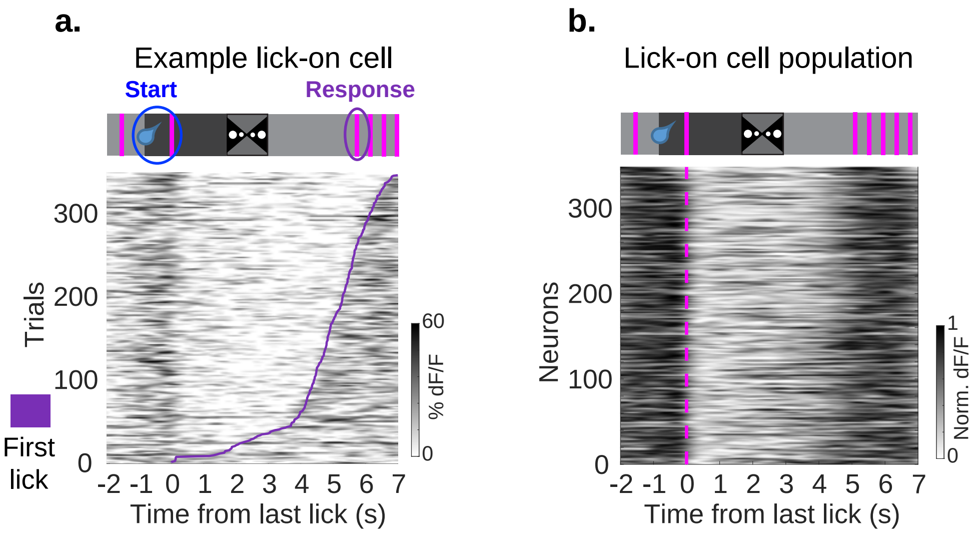


**Supplemental Figure 5:** Lick-on cells

a. Top: schematic of task. Bottom: example lick-on cell. Each row is one trial. Trials are ordered based on the animal's first lick time (purple line). This neuron increases its activity to the first lick and has sustained activity until the last lick.
b. Top: schematic of task. Bottom: all lick-on cells. Neurons are ordered based on the strength of their response to the last lick ($dFF_{aft}/dFF_{bef}$) (see Methods).

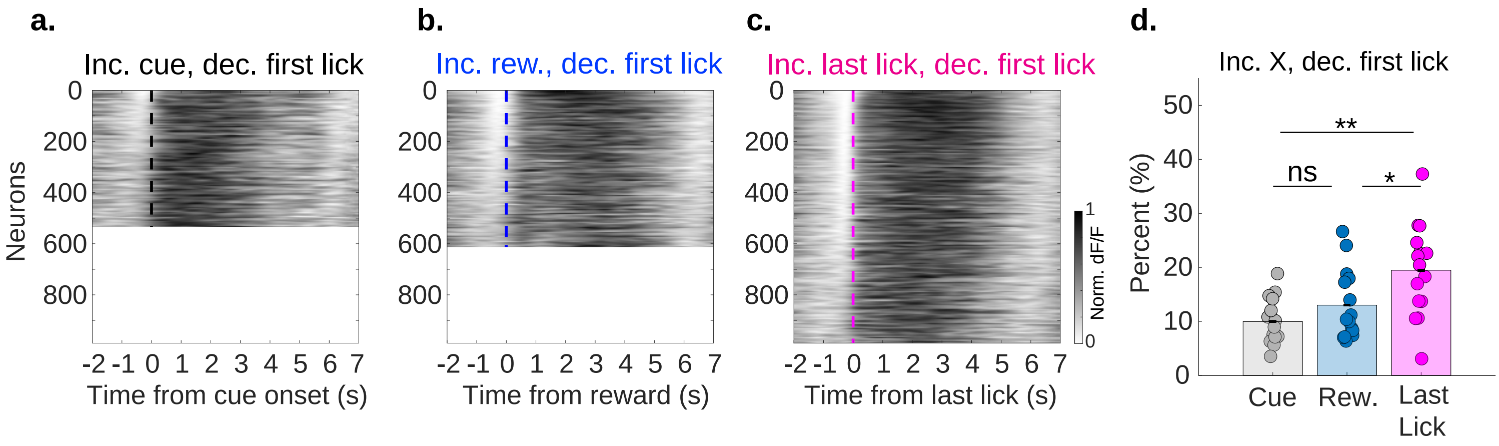


**Supplemental Figure 6:** The percentage of persistently active neurons is higher when aligned to the last lick than aligned to the cue or reward

a. Neurons that increase activity at the cue onset and decrease activity at the first lick. Activity aligned to cue onset (534/5,161 neurons).
b. Neurons that increase activity at the reward delivery and decrease activity at the first lick. Activity aligned to reward delivery (613/5,161 neurons).
c. Neurons that increase activity at the last lick and decrease activity at the first lick (persistently active cells). Activity aligned to last lick (989/5,161 neurons).
d. Percentage of neurons in each of the conditions in S6a-c (cue inc., lick dec.: 9.977 ± 1.109%; rew. inc., lick dec.: 13.011 ± 1.676%; last lick inc., lick dec.: 19.474 ± 2.191%; cue vs. reward: $p = 0.272$, cue vs. last lick: $p = 0.002$, last lick vs. reward: $p = 0.025$; Wilcoxon rank-sum test; 11 animals, 15 sessions).

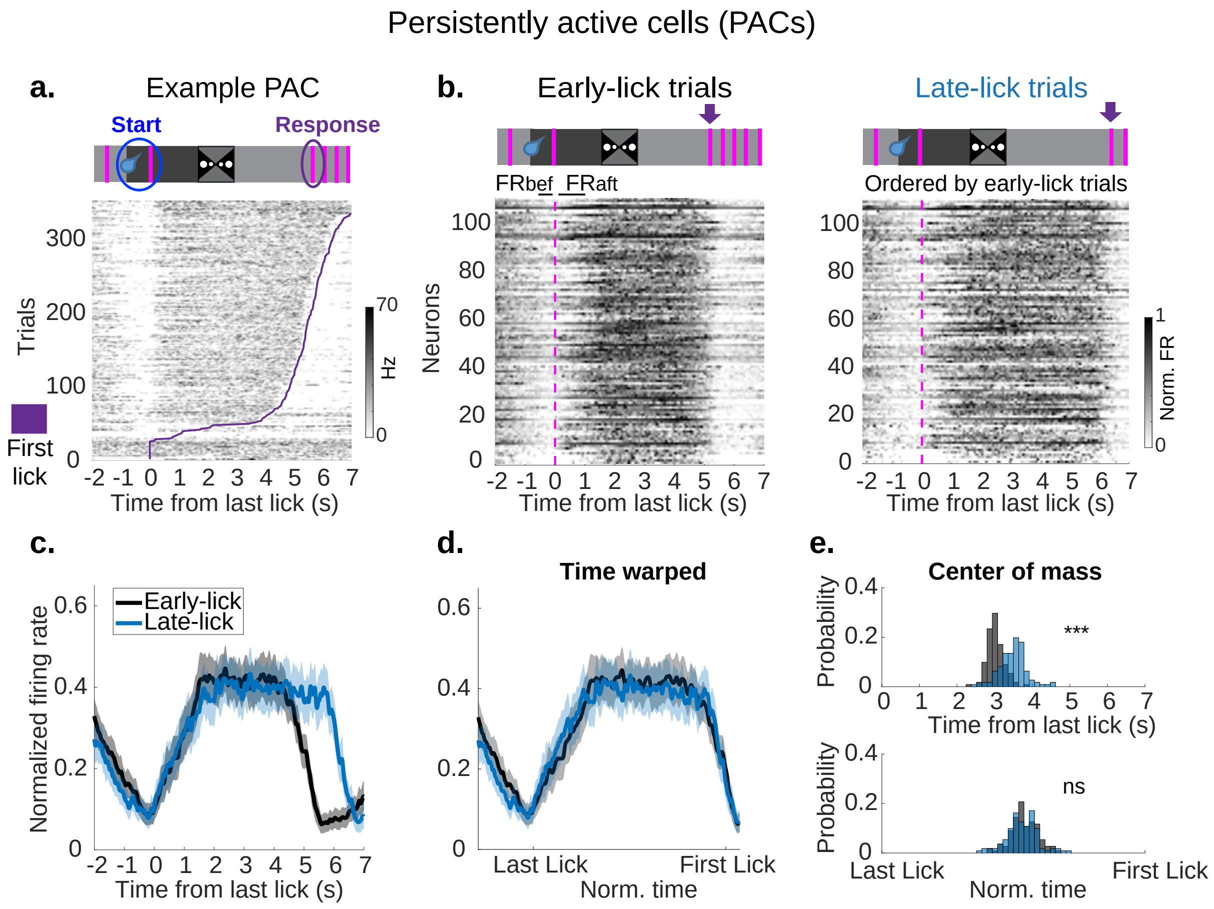


**Supplemental Figure 7:** PACs are present in electrophysiology data

a. Top: schematic of task. Bottom: example persistently active cell. Each row is one trial. Trials are ordered based on the animal's first lick time (purple line).
b. Top left and right: schematic of task. Purple arrow points to the first lick. Bottom left: all persistently active cells on trials where the animals' first lick time is between 4.5-5.5s ("early-lick trials"). Neurons are ordered based on the strength of their response to the last lick (FRaft/FRbef) (see Methods). Bottom right: same neurons in the same order but for trials where the animals' first lick time is 6-7s ("late-lick trials").
c. Averaged normalized firing rate traces for the neurons in S7b. Early-lick trials (black), late-lick trials (blue).
d. Time warped traces from the last lick to the first lick for the data in S7c.
e. Histogram of the center of mass (COM) of persistently active cells. Top: non-warped data. Bottom: warped data (non-warped data COM: early-lick trials: 3.025 ± 0.057s; late-lick trials: 3.485 ± 0.097s; p = 7.203e-21, Kolmogorov–Smirnov test, mean Difference = 0.461, Cohen's D = 1.490; warped data: early-lick trials, 0.542 ± 0.013; late-lick trials, 0.535 ± 0.016; p = 0.512, Kolmogorov–Smirnov test, mean difference = -0.008, Cohen's D = -0.140; 111/952 neurons, 5 animals, 15 sessions).

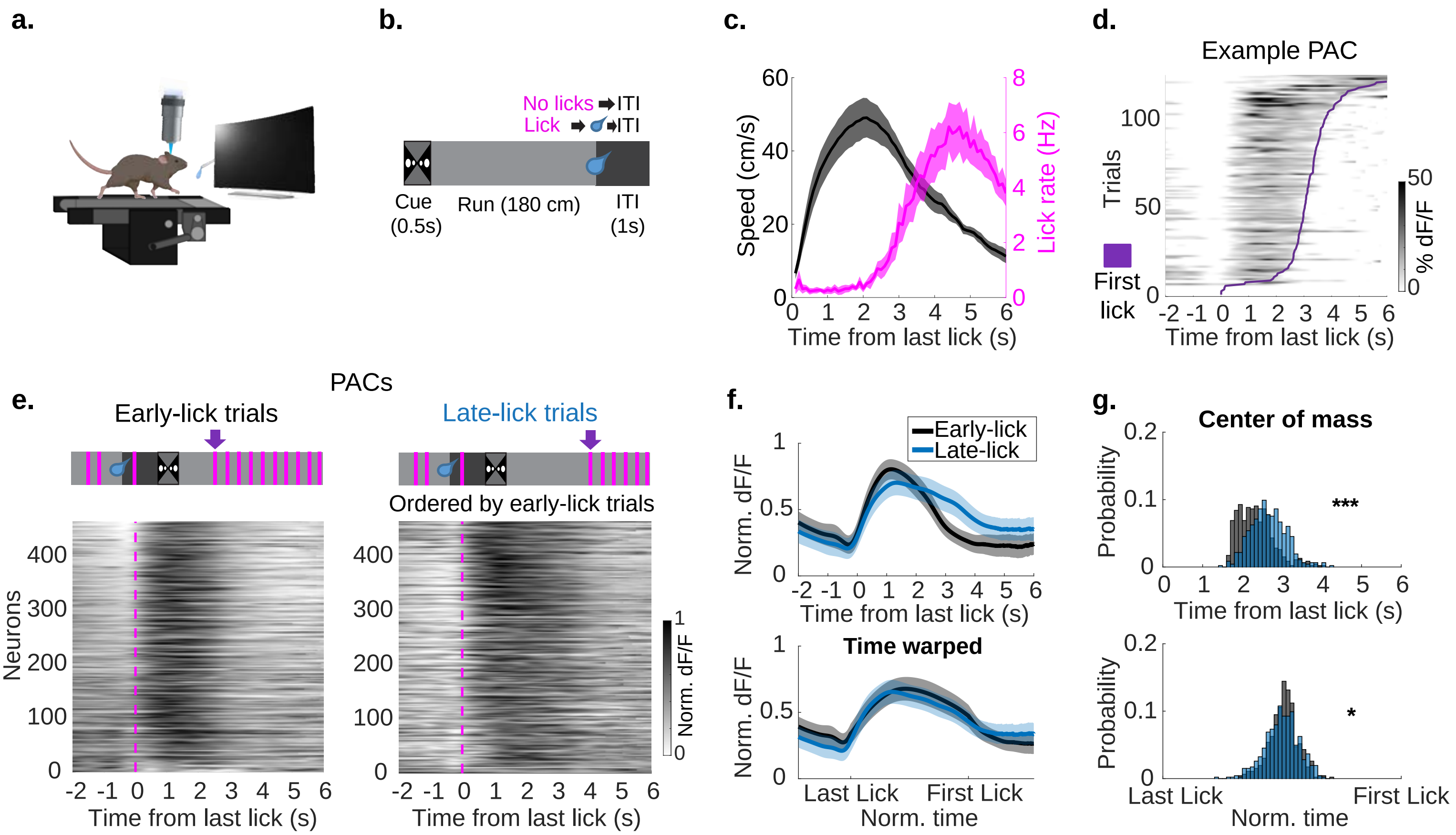


**Supplemental Figure 8:** Persistently active cells generalize to a running task

a. Schematic of running task behavioral apparatus. Head-fixed mice run on a treadmill in front of virtual reality screens and a lick port.
b. Schematic of running task structure. There was a 0.5s visual cue at the trial onset. Then mice had to run 180 cm. After running 180 cm mice had to lick within a 40 cm unmarked reward zone to trigger water reward. Failure to lick within the reward zone resulted in no reward for that trial.
c. Averaged speed trace (black) and licking trace (magenta) for mice performing the running task. The shaded area represents SEM.
d. Example persistently active cell. Each row is one trial. Trials are ordered based on the animal's first lick time (purple line).
e. Top left and right: schematic of task. Purple arrow points to the first lick. Bottom left: all persistently active cells on trials where the animals' first lick time is between 2-3s ("early-lick trials"). Neurons are ordered based on the strength of their response to the last lick ($dFF_{aft}/dFF_{bef}$). Bottom right: same neurons in the same order but for trials where the animals' first lick time is 3.5-4.5s ("late-lick trials").
f. Top: Averaged normalized dF/F traces for the neurons in S8e. Early-lick trials (black), late-lick trials (blue). Bottom: Time warped traces from the last lick to the first lick for the data in S8f, top.
g. Histogram of the center of mass (COM) for persistently active cells. Top: non-warped data. Bottom: warped data (non-warped data COM: early-lick trials: 2.345 ± 0.171s; late-lick trials: 2.658 ± 0.169s; p = 7.731e-22, Kolmogorov–Smirnov test, mean difference: 0.313, Cohen's D: 0.695; warped data COM: early-lick trials: 0.508 ± 0.023; late-lick trials: 0.499 ± 0.027; p = 0.016, Kolmogorov–Smirnov test, mean difference: -0.009, Cohen's D: -0.132;). Although the statistics are significant for both warped and non-warped data, the effect size is small for warped data.

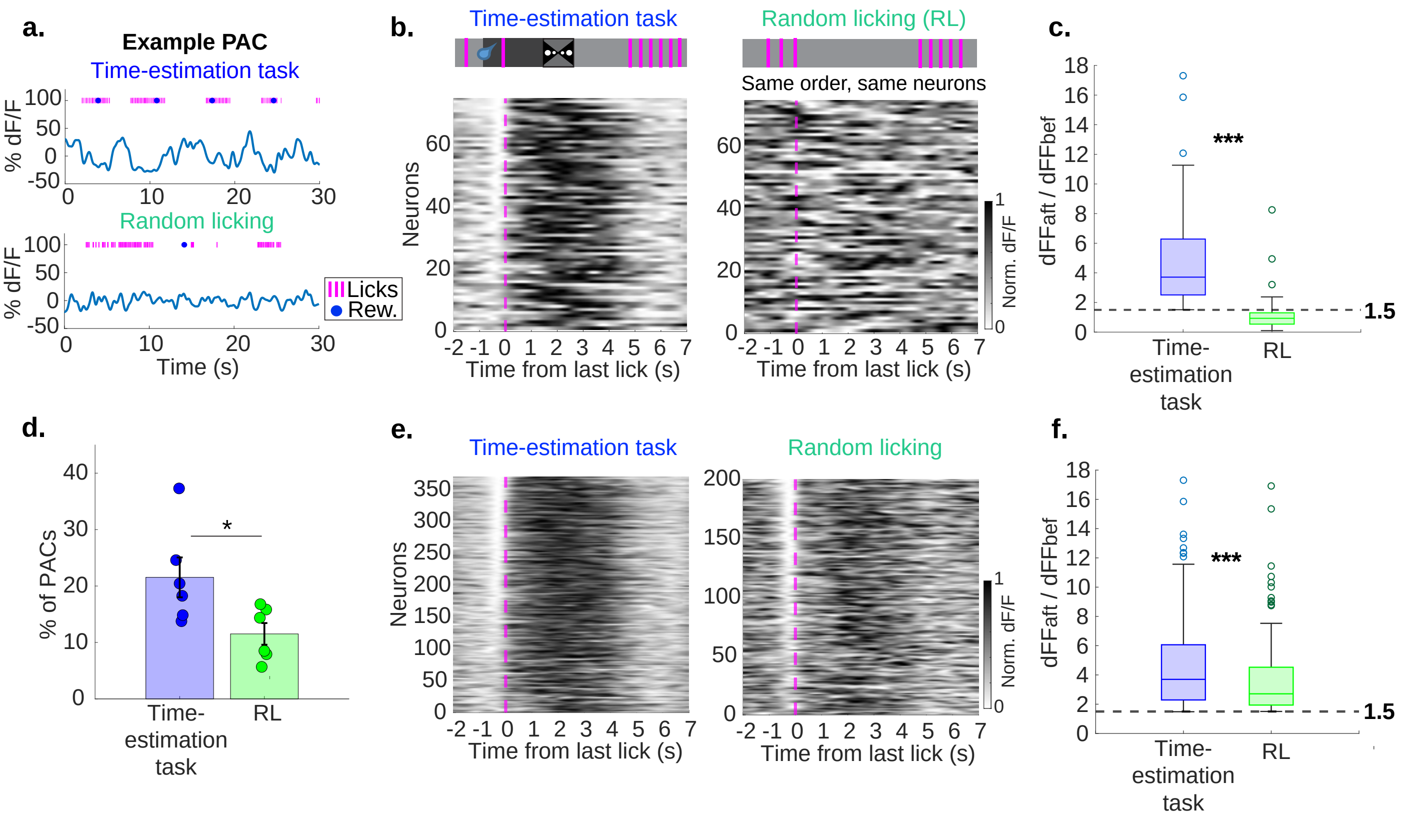


**Supplemental Figure 9:** PACs lose their tuning during random licking

a. Example timeseries data of a PAC during the time-estimation task (top) and during the random licking (RL) session (bottom).

b. Top left: schematic of the time-estimation task. Bottom left: all PACs during the time-estimation task that were also active during the RL session. Neurons are ordered based on the strength of their response to the last lick (dFFaft/dFFbef). Top right: schematic of the RL session. Bottom right: same cells in the same order as bottom left but during the RL session.

c. dFFaft/dFFbef ratio aligned to last lick for neurons in S9b (time-estimation task: 4.725 ± 1.320; RL session: 1.102 ± 0.450; p = 7.589e-14, paired t-test; 75 neurons, 4 animals, 6 sessions).

d. Percentage of PACs identified across all the pyramidal neurons in the time-estimation task or the RL sessions (time-estimation task: 21.533 ± 3.534%; RL sessions: 11.515 ± 1.921%; p = 0.047, paired t-test; 4 animals, 6 sessions).

e. Heatmap of all PACs during the time-estimation task (left), and all PACs during the RL sessions (right). Neurons are ordered based on the strength of their response to the last lick (dFFaft/dFFbef).

f. dFFaft/dFFbef ratio aligned to last lick for neurons in S9e (time-estimation task: 4.490 ± 1.167; RL session: 3.601 ± 0.978; p = 2.737e-05, Wilcoxon rank-sum test; time-estimation task: 370/1,625 neurons; RL session: 200/1,594 neurons, 4 animals, 6 sessions).

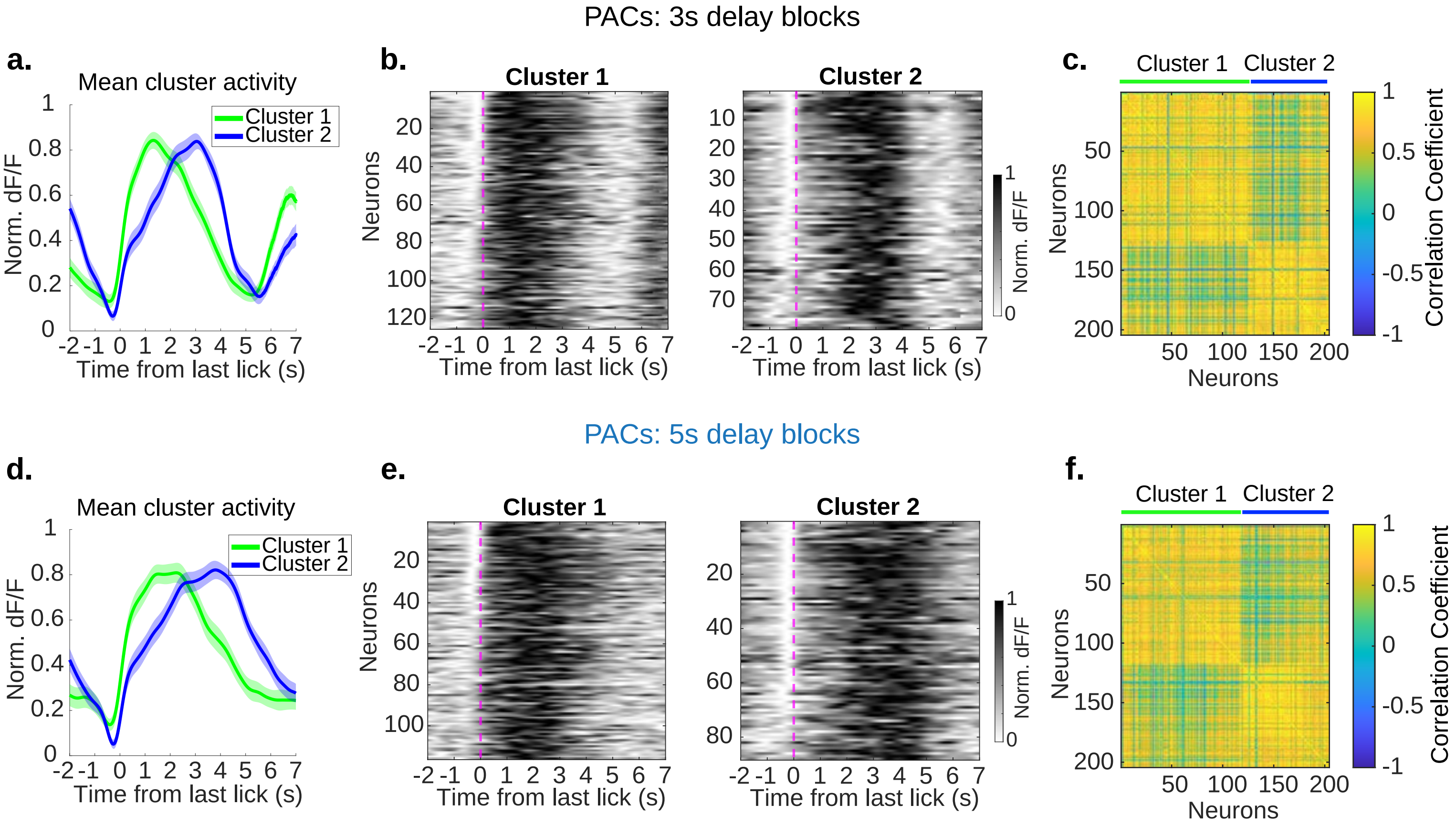


**Supplemental Figure 10:** PACs during the block delay experiment exhibit ramping activity

a. Results from K-means clustering. Averaged normalized dF/F traces from neurons in cluster 1 (green) and cluster 2 (blue) for 3s delay blocks. Neurons in cluster 1 “ramped up” and neurons in cluster 2 “ramped down”.

b. Left: heatmap of cluster 1 neurons sorted by the ratio of their response to last lick ($dFF_{aft}/dFF_{bef}$). Right: Same but for cluster 2 neurons.

c. Correlation matrix of neurons grouped by cluster identity for 3s delay blocks (Cluster 1: 125/204 neurons; Cluster 2: 79/204 neurons).

d-f. Same as a-c but for 5s delay blocks (Cluster 1: 116/204 neurons; Cluster 2: 88/204 neurons).

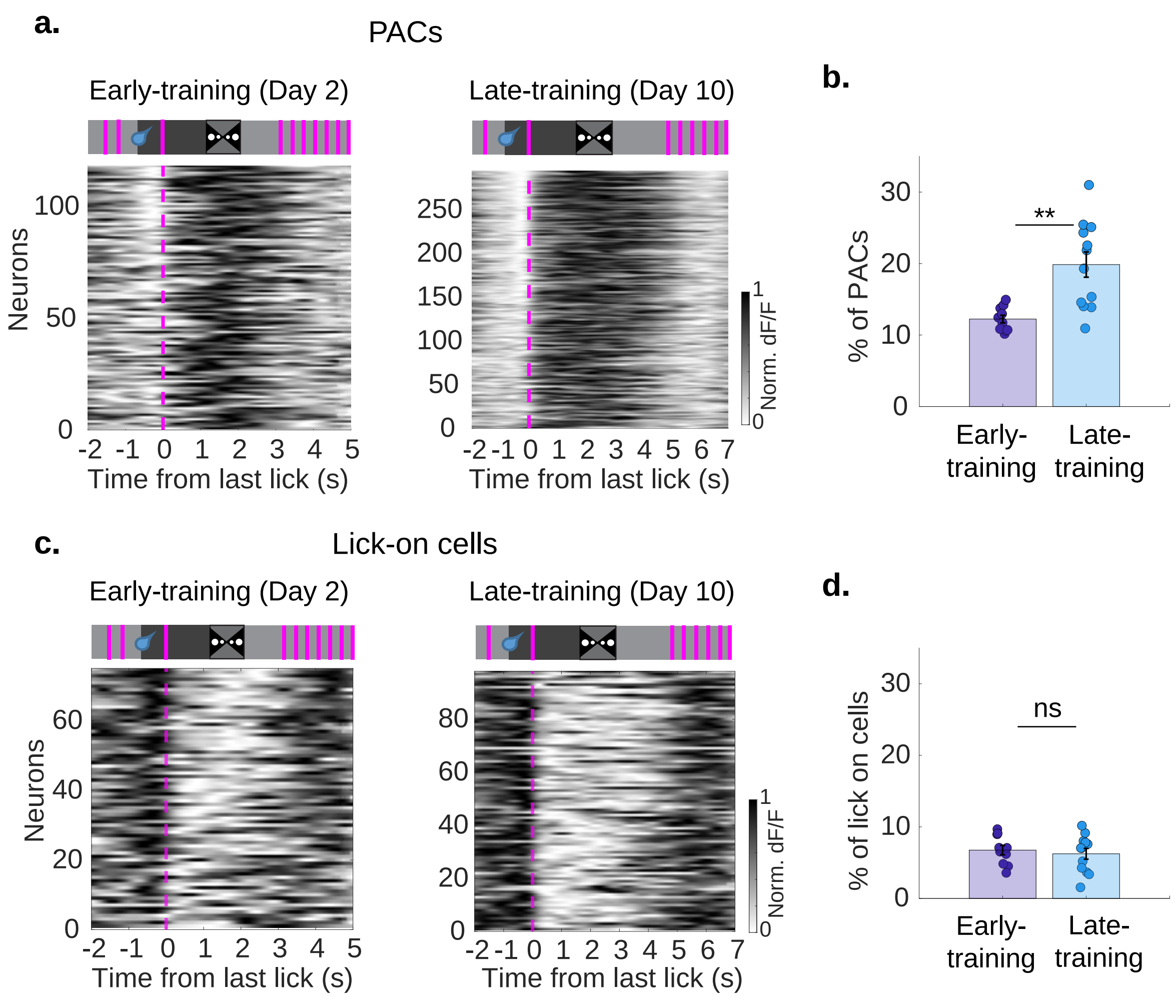


**Supplemental Figure 11:** PAC percentage increases with learning

a. Data from trials with clear first and last licks. Left: all PACs across 5 animals on early-training day 2. Right: all PACs from the same animals but on late-training day 10.
b. Percentage of PACs increases from early- to late-training (early-training: 12.237 ± 0.531%; late-training: 19.859 ± 1.767%; p = 0.001, Wilcoxon rank-sum test; 4 animals, early-training: 10 sessions, late-training: 12 sessions).
c. Data from trials with clear first and last licks. Left: all lick-on cells across 5 animals on early-training day 2. Right: all lick-on cells from the same animals but on late-training day 10.
d. Percentage of lick-on cells does not increase from early- to late-training (early-training: 6.755 ± 0.654%; late-training: 6.238 ± 0.752%; p = 0.767, Wilcoxon rank-sum test).
